# GAS6 and AXL promote insulin resistance by rewiring insulin signaling and increasing insulin receptor trafficking to endosomes

**DOI:** 10.1101/2023.10.19.563096

**Authors:** Céline Schott, Amélie Germain, Julie Lacombe, Monica Pata, Denis Faubert, Jonathan Boulais, Peter Carmeliet, Jean-François Côté, Mathieu Ferron

## Abstract

Growth-arrest specific 6 (GAS6) is a secreted protein that acts as a ligand for TAM receptors (TYRO3, AXL and MERTK). In humans, GAS6 circulating levels and genetic variations in *GAS6* are associated with hyperglycemia and increased risk of type 2 diabetes. However, the mechanisms by which GAS6 influences glucose metabolism are not understood. Here, we show that *Gas6* deficiency in mice increases insulin sensitivity and protects from diet-induced insulin resistance. Conversely, increasing GAS6 circulating levels is sufficient to reduce insulin sensitivity *in vivo*. GAS6 inhibits the activation of the insulin receptor (IR) and reduces insulin response in muscle cells in vitro and in vivo. Mechanistically, AXL and IR fom a complex, while GAS6 reprograms signaling pathways downstream of IR. This results in increased IR endocytosis following insulin treatment. This study contributes to a better understanding of the cellular and molecular mechanisms by which GAS6 and AXL influence insulin sensitivity.

**ARTICLE HIGHLIGHTS:**

- GAS6 deficient mice are characterized by improved glucose tolerance, increased insulin sensitivity and are protected from diet-induced insulin resistance.
- The GAS6 receptor AXL is expressed in muscle cells and forms a complex with the insulin receptor.
- GAS6 reprograms insulin signaling by modulating the phosphorylation of proteins implicated in endocytosis, vesicle-mediated transport, and membrane trafficking.
- AXL and GAS6 promotes insulin receptor endocytosis following insulin treatment.

## INTRODUCTION

At the root of the pathophysiology of type 2 diabetes (T2D) lies insulin resistance, an insufficient response to insulin in skeletal muscle, white adipose tissue, and liver. In normal conditions, insulin mediates its actions by binding to the insulin receptor (IR), triggering IR autophosphorylation of its tyrosine kinase domain. These molecular events result in the recruitment and phosphorylation of insulin receptor substrates (IRS) proteins leading to the activation of the phosphatidylinositol 3-kinase (PI3K)/protein kinase B (AKT) pathway that plays a critical role in the molecular response to insulin (1). Alterations in IR and PI3K/AKT activation or phosphorylation in response to insulin impact several cellular pathways including glucose uptake (2). In addition, a reduction in the level of activation of the insulin signaling cascade in muscle has been associated with insulin resistance and contributes to the development of T2D (3). However, the cellular mechanisms involved in the development of muscle insulin resistance are still not fully understood.

Growth-arrest specific 6 (GAS6) is a secreted protein initially identified as upregulated in growth-arrested fibroblasts (4). GAS6 is composed of an N-terminal Gla domain containing γ- carboxyglutamic acids, four central epidermal growth factor-like domains and two laminin G-like (LG) domains (5). GAS6 functions as a ligand for the TAM family of receptor tyrosine kinases (RTK), which includes 3 members: TYRO3, AXL and MERTK (6). Protein S (PROS), which functions as an anticoagulant factor, shares 44% of its amino acid sequence with GAS6. However, GAS6 and PROS may have different functions *in vivo* given that mice lacking *Pros1* dies in utero from hemorrhages, while *Gas6^-/-^*mice are viable and do not suffer from spontaneous bleeding or thrombosis (7; 8). Recent studies have established that only GAS6 can activate AXL, while TYRO3 and MERTK act as both PROS and GAS6 receptors (9; 10), suggesting that the non-redundant functions of GAS6 are mediated by AXL. TAM receptors have been implicated in several homeostatic functions in adult mice including spermatogenesis, clearance of apoptotic cells, suppression of inflammation, and platelet activation (11–14), but are also involved in tumor progression and metastasis (15; 16). Nevertheless, the physiological functions of GAS6 remains unclear, since mice lacking *Gas6* are apparently devoid of phenotype under normal conditions (8; 17).

Studies in humans suggest a potential role for GAS6 and AXL in insulin sensitivity, glucose homeostasis and T2D (18–20). Serum levels of GAS6 were found to be elevated in obese adolescents compared to lean subjects and circulating levels of GAS6 were strongly correlated with insulin resistance in this population (20). Genome-wide association studies (GWAS) have identified a positive association between non-coding single-nucleotide polymorphisms (SNP) in the human *GAS6* gene and the circulating levels of glycated hemoglobin (HbA1c), a marker of hyperglycemia (21; 22). Although these data suggest that GAS6 may play a role in the development of insulin resistance and T2D in humans, the physiological and molecular mechanisms through which GAS6 and its receptor(s) might influence these metabolic disorders are currently unknown.

Here, we first show that the absence of GAS6 in mice improves insulin sensitivity, without affecting insulin secretion, body weight or energy expenditure. Next, we establish that AXL forms a complex with IR in an insulin-dependent manner and, through quantitative phosphoproteomics, we demonstrate that GAS6 profoundly modifies insulin signaling in muscle cells leading to increased insulin receptor internalization. This study uncovers a crosstalk between GAS6 and insulin signaling and provide the basis of the molecular mechanism by which GAS6 promotes insulin resistance and T2D.

## RESEARCH DESIGN AND METHODS

### Mice

*Gas6^-/-^* mice on a C57BL/6J genetic background and on a FVB/N genetic background were previously described (8; 16). *ApoE-Gas6^Tg^* transgenic mice were generated in our laboratory as described in Supplemental Material. All strains were maintained in an IRCM specific pathogen-free animal facility under 12-hour dark/12-hour light cycles. Mice were fed ad libitum a normal chow diet (Teklad global 18% protein rodent diet; 2918; Envigo), unless otherwise specified. Male mice were used in all experiments.

### Cell Lines

The C2C12, HEK293, HEK293 Flp-In T-Rex cells expressing or not AXL-BirA*-Flag and L6-GLUT4myc cells are described in Supplemental Material.

### Metabolic Analysis

For glucose tolerance tests (GTT), mice were fasted for 16 hours and glucose (2g/kg or 1.5g/kg) was administered intraperitoneally (i.p.) or orally. For insulin tolerance test (ITT), mice were fasted 6 hours, injected i.p. with recombinant insulin (Humulin R Lilly) (0.6 U/kg for *Gas6^-/-^* mice or 0.75 U/kg for *ApoE-Gas6^Tg^*transgenic mice). For pyruvate tolerance test, mice were fasted 16 hours and injected i.p. with sodium pyruvate (2 g/kg). Blood glucose levels were measured using a Bravo glucometer (EndoMedical Inc.). In vivo glucose stimulated insulin secretion test (GSIS) were performed after 16 hours of fasting and i.p. injection with 3 g/kg glucose. Serum insulin was measured with a commercial insulin ELISA kit (Mercodia).

Mouse physiological parameters were measured using an 8-chamber Promethion Continuous Metabolic System (Sable Systems International) as we previously reported (23). Body composition was evaluated with a body composition analyzer (echo-MRI). When indicated, mice were fed a 60% high-fat high-sucrose diet (D12331; Research Diets) for 12 weeks starting from 5 weeks of age and fasted for 6 hours for GTT (1 g/kg glucose) and ITT (0.75 U/kg insulin).

### DNA Constructs

All constructs were generated by PCR using primers included in **Supplemental Table 1** and as described in Supplemental Material.

### Production and Purification of Recombinant GAS6

Carboxylated and uncarboxylated mouse Gas6 production, purification and quantification were performed as described in Supplemental Material.

### Treatment with GAS6

For 15-minute GAS6 stimulation, C2C12 cells were starved overnight in DMEM media with 0.5% FBS. After two PBS 1X washes, cells were cultured for 2h in serum-free medium supplemented with 10 mM of HEPES pH 7.4 and 0.1% BSA and stimulated for 15 minutes with 200 ng/ml of recombinant carboxylated or uncarboxylated GAS6. For 8 and 24-hour GAS6 treatment, C2C12 and L6-GLUT4myc cells were incubated in serum-free medium with or without recombinant GAS6 (200ng/ml).

### Co-Immunoprecipitation and Western Blot

HEK293 cells were transfected with pcDNA5-AXL-BirA*-Flag and pcDNA3-IR-HA and processed as described in Supplemental Material. For in vivo insulin signaling analysis, mice were fasted 16 hours and anesthetized with a drug mixture of ketamine hydrochloride and xylazine before being injected with saline or insulin (0.5U/kg) in the inferior vena cava. Mice were sacrificed via cervical dislocation after 12 minutes and epididymal WAT, liver and gastrocnemius muscles were collected and analyzed by Western blot. Antibodies used for western blot are listed in **Supplemental Table 2**.

### Streptavidin Pull-Down

HEK293 Flp-In T-Rex cells expressing AXL-BirA*-Flag were cultured during 24 hours with biotin (50µM) and tetracycline (1µg/ml), then cultured in 0.5% FBS DMEM overnight and incubated in serum-free medium (with 0.1% BSA and 10 mM HEPES) for 2 hours followed by 15-minute stimulation with insulin (100 nM). Streptavidin pull-down was performed as described in Supplemental Material.

### Immunofluorescence Confocal Microscopy

HEK293 cells were transfected with pcDNA5-AXL-BirA*-Flag and/or pmScarlet-C1-Rab7 and processed as described in Supplemental Material. Antibodies used for immunofluorescence are listed in **Supplemental Table 2**. Images were acquired with Zeiss LSM700 confocal microscope with a 63X oil objective. We used Image J software to quantify Insulin Receptor (IR) and Rab7 localization. For each different condition, IR signal intensity (Alexa488) was measured across the cell from the nucleus to the plasma membrane and normalized to cell size (n=30 cells per condition, from 3 independent experiments). Colocalization percentage was measured as IR and Rab7 overlapping area for each cell over cell area. Staining and imaging of IR in skeletal muscle sections was performed and quantified as described in Supplemental Material.

### Gene Expression

Tissue and cell total RNA extraction and isolation were performed as described in Supplemental Material.

### Protein Digestion for Phosphoproteome Analysis

C2C12 cells were incubated in serum-free medium with or without recombinant GAS6 (200ng/ml) for 24 hours and treated or not with insulin (100 nM) for 15 minutes. Protein extracts were digested with benzonase as described in Supplemental Material.

### Isobaric Peptide Labeling and LC-MS/MS Analysis

Protein digests were labelled using the 6-plex tandem mass tag reagent (Thermo Fisher Scientific) and analyzed by LC-MS/MS as described in Supplemental Material.

### Bioinformatics

We considered as modulated phosphosites those showing a normalized Log2 (≤L−0.5 or ≥0.5) between two given conditions and *P* < 0.05. Detailed analysis of the phophopeptides is available in Supplemental Tables 3-6: https://figshare.com/s/43f5bf99204c1d928e05. Reactome pathway analyses were performed to functionally annotate proteins using Gene Set Enrichment Analysis (GSEA) with an FDR threshold of 0.05. Graphical network representations of protein-protein interaction were performed with the STRING app (11.5) in Cytoscape (3.9.1).

### Statistics

Statistical analyses were performed using GraphPad Prism software as described in Supplemental Material.

### Data availability statements

The datasets generated during and/or analyzed during the current study are available from the corresponding author upon reasonable request.

### Resource availability statements

The *ApoE-Gas6^Tg^* transgenic mouse line generated and analyzed during the current study is available from the corresponding author upon reasonable request. The pcDNA3-IR-HA, pcDNA5-AXL-BirA*-Flag and pmScarlet-C1-Rab7 DNA constructs, the AXL-BirA*-Flag and mGAS6-Myc-6xHis HEK293 Flp-In T-Rex cell lines, and the phospho-AXL (Y702) antibody generated during the current study are available from the corresponding author upon request.

## RESULTS

### GAS6 deficiency improves glucose tolerance and insulin sensitivity in mice fed a normal chow diet

To determine GAS6 function in glucose homeostasis, we characterized the metabolic phenotype of 3-month-old *Gas6^-/-^* male mice fed a standard chow diet on two inbred genetic backgrounds: FVB/N and C57BL/6J. These two strains were selected because they are both widely used in metabolic studies but have significant differences in whole-body glucose metabolism (24). When compared to control littermates on the same genetic background, *Gas6^-/-^* mice did not display any differences in energy expenditure parameters, physical activity, body weight and food intake (**Fig.S1A-P**). However, on both genetic backgrounds, *Gas6^-/-^*mice have significantly reduced serum glucose levels in fed conditions (**Fig. 1A**-**B**) or after a glucose load as assessed through intraperitoneal and oral glucose tolerance tests (GTTs) (**Fig. 1C-D** and **Fig.S1Q**).

**Figure 1.**
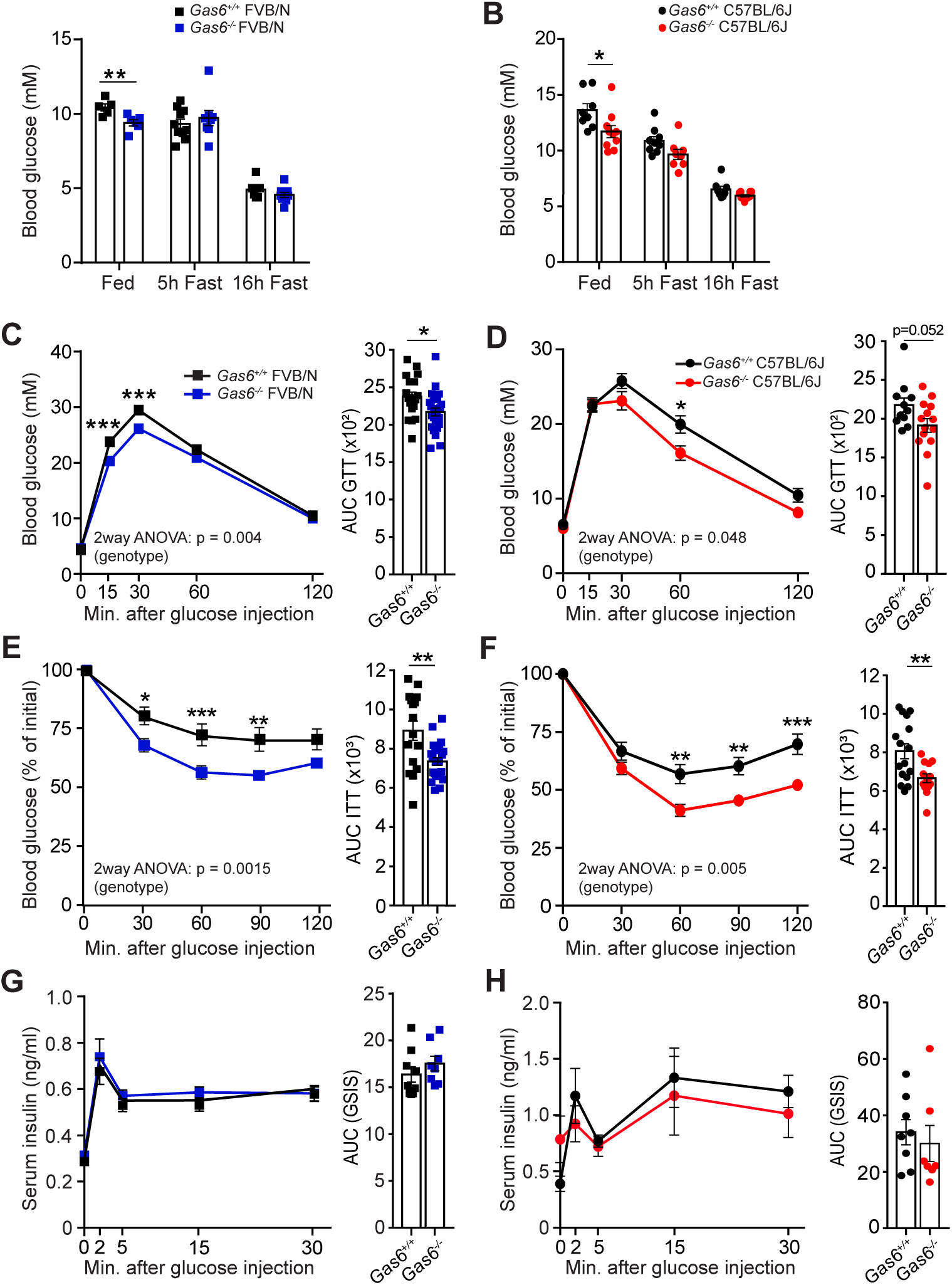
GAS6 deficiency improves glucose tolerance and insulin sensitivity in FVB/N and C57BL/6J mice under chow diet. Metabolic parameters of 3-month-old *Gas6^+/+^* and *Gas6^-/-^* FVB/N (blue) and C57BL/6J (red) male mice. **(A-B)** Fed and fasting (5h or 16h) blood glucose levels (n=5-10). **(C-D)** Intraperitoneal glucose tolerance Test (IPGTT) and area under the curve (AUC) (n=11-25). Mice were fasted for 16 hours and injected i.p. with 2g/kg of glucose. **(E-F)** For insulin tolerance test (ITT), mice were fasted for 5 hours and injected i.p. with 0.6 U/kg of insulin (n=14-25). Results are expressed as a percentage of basal glycemia and AUC. **(G-H)** Glucose stimulated insulin secretion (GSIS) test and AUC in mice injected i.p. with 3g/kg of glucose after 16 hours of fasting (n=7-9). Results represent mean ± SEM, \**P* < 0.05, \*\**P* < 0.01, \*\*\**P* < 0.001 by unpaired, 2-tailed Student’s *t* test **(A-B** and for AUC in **C-H)**, or by two-way ANOVA for repeated measurements with Bonferroni’s post tests **(C-H)**.

Insulin tolerance tests (ITT) suggested that *Gas6^-/-^* mice on FVB/N or C57BL/6J backgrounds have improved insulin sensitivity (**Fig. 1E-F**). Glucose-stimulated insulin secretion tests (GSIS) showed that *Gas6^-/-^*mice had normal insulin secretion in response to glucose (**Fig. 1G-H**). Gluconeogenesis from pyruvate was not significantly affected in *Gas6^-/-^* mice, as assessed by pyruvate tolerance tests (PTT) (**Fig.S1R-S**). These data suggest that GAS6-deficiency improves glucose tolerance by affecting insulin sensitivity, but not insulin secretion or gluconeogenesis.

### *Gas6*-deficient mice are protected from diet-induced insulin resistance

We next investigated whether GAS6 deficiency could have beneficial effects on glucose metabolism in a model of obesity induced by a high-fat and high-sucrose diet (HFHS). This study was performed only in mice on C57BL/6J genetic background, since this inbred strain is more susceptible to diet-induced obesity (25). Twelve weeks of HFHS feeding equally affected body weight, body composition (fat and lean mass) and the weight of epididymal and subcutaneous white adipose tissues (eWAT and scWAT) in *Gas6^+/+^* and *Gas6^-/-^* mice (**Fig.2A-E**). Obesity is associated with WAT inflammation characterized by an increased number of activated macrophages that contribute to insulin resistance in this tissue (26) and TAM receptors have been previously implicated in immune cell functions (27). However, although the number and area of F4/80 positive macrophages in WAT following the HFHS diet was increased as expected, no significant impact of the genotype was observed (**Fig.S2A-B**).

**Figure 2.**
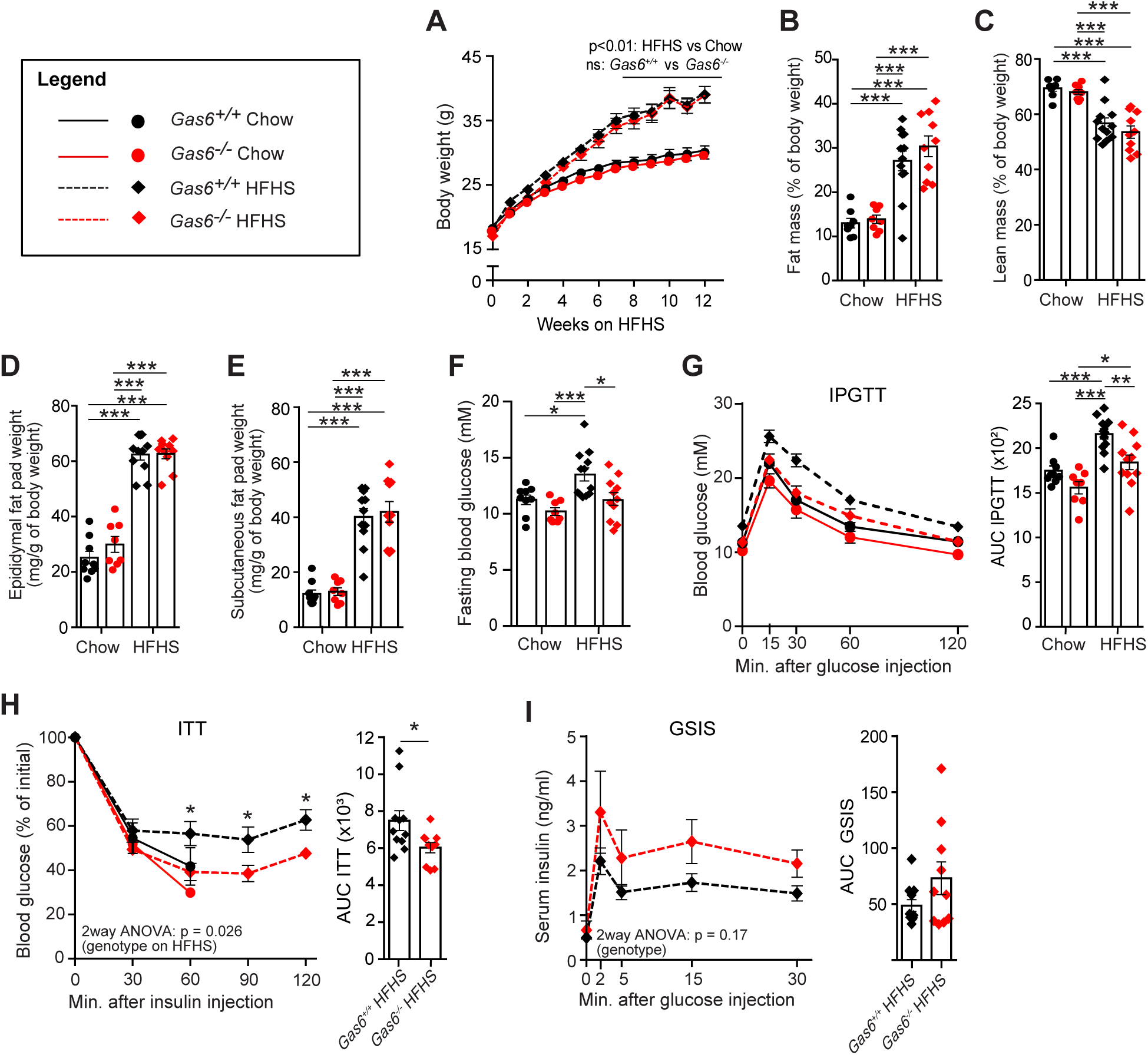
GAS6 deficiency protects from High-Fat/High-Sucrose diet-induced insulin resistance independently of body weight. **(A)** Body weight, percentage of total **(B)** fat and **(C)** lean mass over body weight and **(D)** epididymal and **(E)** subcutaneous fat pad weight of *Gas6^+/+^* and *Gas6^-/-^* C57BL/6J mice fed a chow diet (Chow) or a High-Fat/High-Sucrose diet (HFHS) for 12 weeks (n=8-12). **(F)** Six hours fasting blood glucose. **(G)** IPGTT results and AUC (n=8-12). Mice were fasted for 5 hours and injected i.p. with 1g/kg glucose. **(H)** ITT in mice after a 5h-fast and injected i.p. with 0.75U/kg of insulin (n=7-11). Results are expressed as a percentage of basal glycemia and as AUC. **(I)** GSIS test in mice fasted for 16 hours and injected i.p with 3g/kg glucose (n=10-11) and AUC. Results represent mean ± SEM. \**P* < 0.05, \*\**P* < 0.01, \*\*\**P* < 0.001, by two-way ANOVA for repeated measurements with Bonferroni’s post tests **(A** and **G-I)**, by one-way ANOVA with by Bonferroni’s post tests **(B-F** and AUC for **G)** or by unpaired, 2-tailed Student’s *t* test (AUC in **H-I)**.

Following a 6h fast, control mice (*Gas6^+/+^*) under HFHS diet displayed a significant increase in blood glucose in comparison with those fed a chow diet (**Fig.2F**). In contrast, fasting glycemia in *Gas6^-/-^* mice did not increase following HFHS diet feeding and remained comparable with control chow-fed mice (**Fig.2F**). Strikingly, glucose tolerance was similar between *Gas6^-/-^*mice fed an HFHS diet and *Gas6^+/+^* mice fed a chow diet, whereas *Gas6^+/+^* mice fed an HFHS diet were significantly more glucose intolerant (**Fig.2G**). Insulin sensitivity was also markedly improved in *Gas6^-/-^* mice fed an HFHS diet compared to *Gas6^+/+^* littermates fed with the same diet (**Fig.2H**). Insulin secretion following a glucose load (GSIS) tended to be higher in some of the *Gas6-* deficient mice under an HFHS diet, but this difference did not reach statistical significance (**Fig.2I**). These results indicate that the inactivation of *Gas6* can prevent insulin resistance and glucose intolerance induced by HFHS diet.

### Increasing GAS6 circulating levels reduces insulin sensitivity in mice

We then aimed to determine if increasing circulating GAS6 levels is sufficient to decrease insulin sensitivity. For that purpose, we generated a transgenic gain-of-function mouse model in which the liver-specific regulatory sequences of the human *APOE* gene (28) were used to achieve ectopic expression of a Myc-6×His-tagged version of mouse GAS6 in hepatocytes and secretion in blood (**Fig.3A**). Overexpression of secreted proteins in hepatocytes using the human *APOE* promoter was previously shown to result in significantly increased circulating levels of these proteins. Moreover, the liver being a major site of production for vitamin K-dependent proteins such as prothrombin, GAS6 ectopically produced by this organ will be fully γ-carboxylated and active.

**Figure 3.**
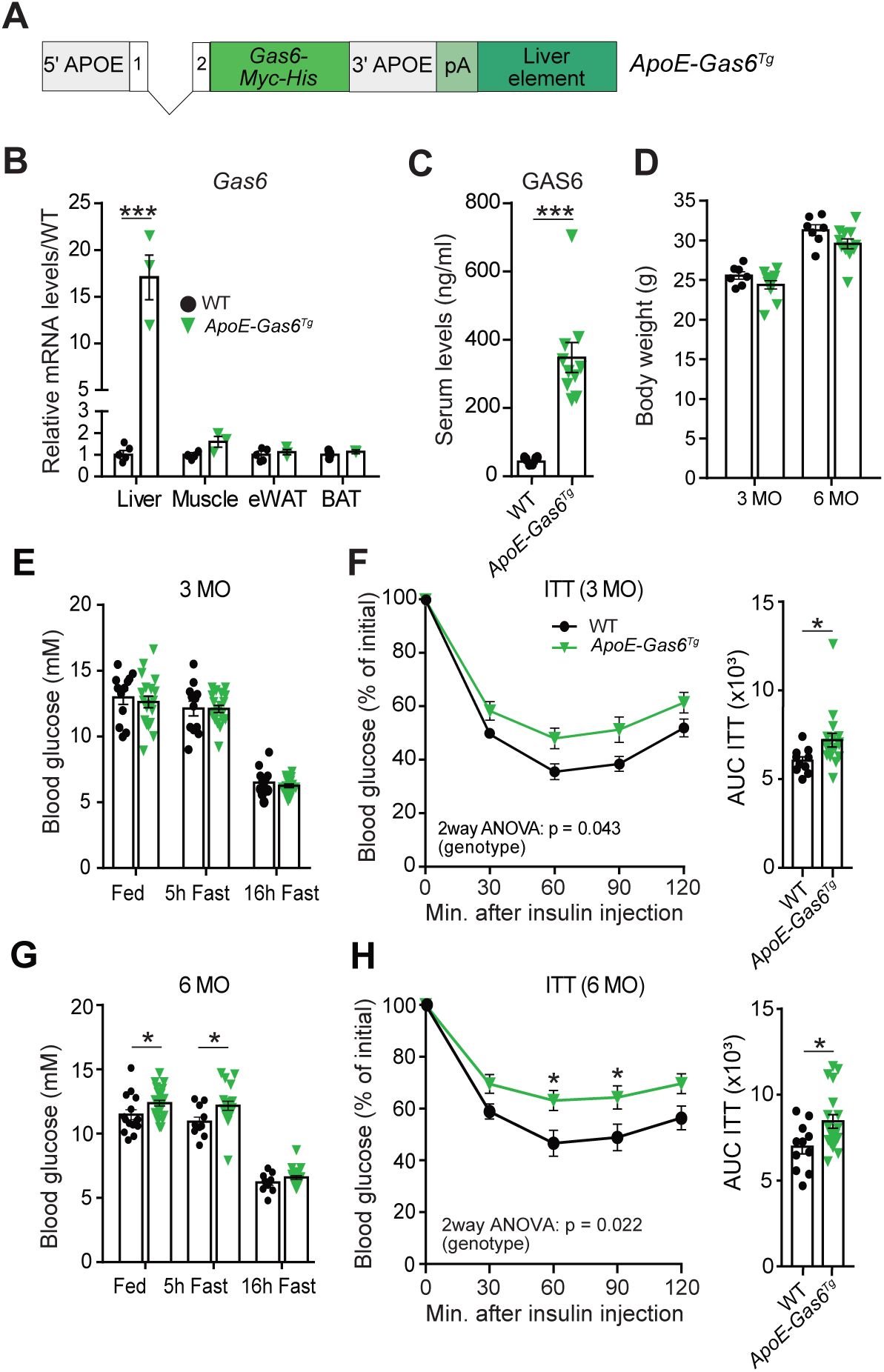
Increasing GAS6 circulating levels reduces insulin sensitivity. **(A)** Schematic representation of the construct used to generate *ApoE-Gas6^Tg^* mice. *Gas6-Myc-6xHis* was cloned in the pLIV.7 vector containing exon 1 and 2, liver-specific regulatory sequences and 5′ and 3′ flanking sequences of the human *APOE* gene. **(B)** *Gas6* gene expression in tissues from WT and *ApoE-Gas6^Tg^* mice was assessed by quantitative PCR (qPCR) and normalized to *Actb* (*n* = 3-5). **(C)** GAS6 serum levels in WT and *ApoE-Gas6^Tg^* mice were measured by ELISA (n=6-10). **(D)** Body weight of WT and *ApoE-Gas6^Tg^*mice at 3 and 6 months of age (n=7-11). **(E** and **G)** Fed and fasting (5h or 16h) blood glucose levels in WT and *ApoE-Gas6^Tg^* mice at 3 and 6 months of age (n=12-24). **(F** and **H)** ITT results at 3 and 6 months of age; mice were fasted for 5 hours and injected i.p. with 0.75U/kg of insulin (n=10-19). Results are expressed as a percentage of basal glycemia and as AUC. Results represent mean ± SEM, \**P* < 0.05, \*\**P* < 0.01, \*\*\**P* < 0.001 by unpaired, 2-tailed Student’s *t* test **(B-E and G** and AUC in **F** and **H)**, or by two-way ANOVA for repeated measurements with Bonferroni’s post tests **(F and H)**.

*ApoE-Gas6^Tg^* mice were generated and maintained on a pure C57BL/6J genetic background. Quantitative PCR (qPCR) confirmed that *Gas6* mRNA levels were ∼18 times higher only in the liver of the *ApoE-Gas6^Tg^* mice compared to wild type (WT) littermate mice (**Fig.3B**). Circulating levels of GAS6, measured using a specific ELISA assay, were approximately 8 times higher in the transgenic animals as compared to WT (**Fig.3C**).

Systemic elevation of GAS6 had no impact on body weight at 3 or 6 months of age compared to WT mice fed a standard chow diet (**Fig.3D**). At 3 months of age, *ApoE-Gas6^Tg^* mice did not show any changes in glycemia, in fed or fasted conditions (**Fig.3E**). However, a significantly reduced insulin sensitivity was detected at the same age in *ApoE-Gas6^Tg^* mice (**Fig.3F**). Six-month-old *ApoE-Gas6^Tg^*mice displayed a significantly increased glycemia in fed conditions and after a 5h fasting period (**Fig. 3G**), and insulin sensitivity was still reduced compared to WT mice (**Fig.3H**). Glucose tolerance was not affected in *ApoE-Gas6^Tg^* mice (**Fig. S3A-B**), suggesting that reduced insulin sensitivity was counterbalanced by glucose disposal mechanisms independently of insulin (29). Therefore, increasing GAS6 circulating levels is sufficient to reduce insulin sensitivity *in vivo*.

### AXL is expressed and activated by GAS6 in muscle cells

The expression pattern of GAS6 and TAM receptors across mouse tissues involved in glucose metabolism was investigated. *Gas6* was strongly expressed in eWAT and scWAT but significantly less in BAT, liver and pancreatic islets (**Fig.4A**). *Gas6* mRNA was also highly expressed in skeletal muscles (**Fig.4A)**. *Axl* and *MerTK*, but not *Tyro3*, were highly expressed in adipose tissues (**Fig.4B**). *MerTK* was more expressed than *Axl* in liver (**Fig.4C**). TAM receptors and GAS6 were expressed at very low level in mouse islets **(Fig.4D**). Interestingly, *Axl* was the most highly expressed TAM receptor in all types of muscles compared to *MerTK* which was weakly present, while *Tyro3* was undetectable (**Fig.4E**). Since *Gas6* and *Axl* are strongly expressed in skeletal muscle and because this tissue is responsible for ∼80% of glucose uptake in response to insulin in the postprandial state (30), we further investigated a potential implication of GAS6 in the response to insulin in skeletal muscle cells.

**Figure 4.**
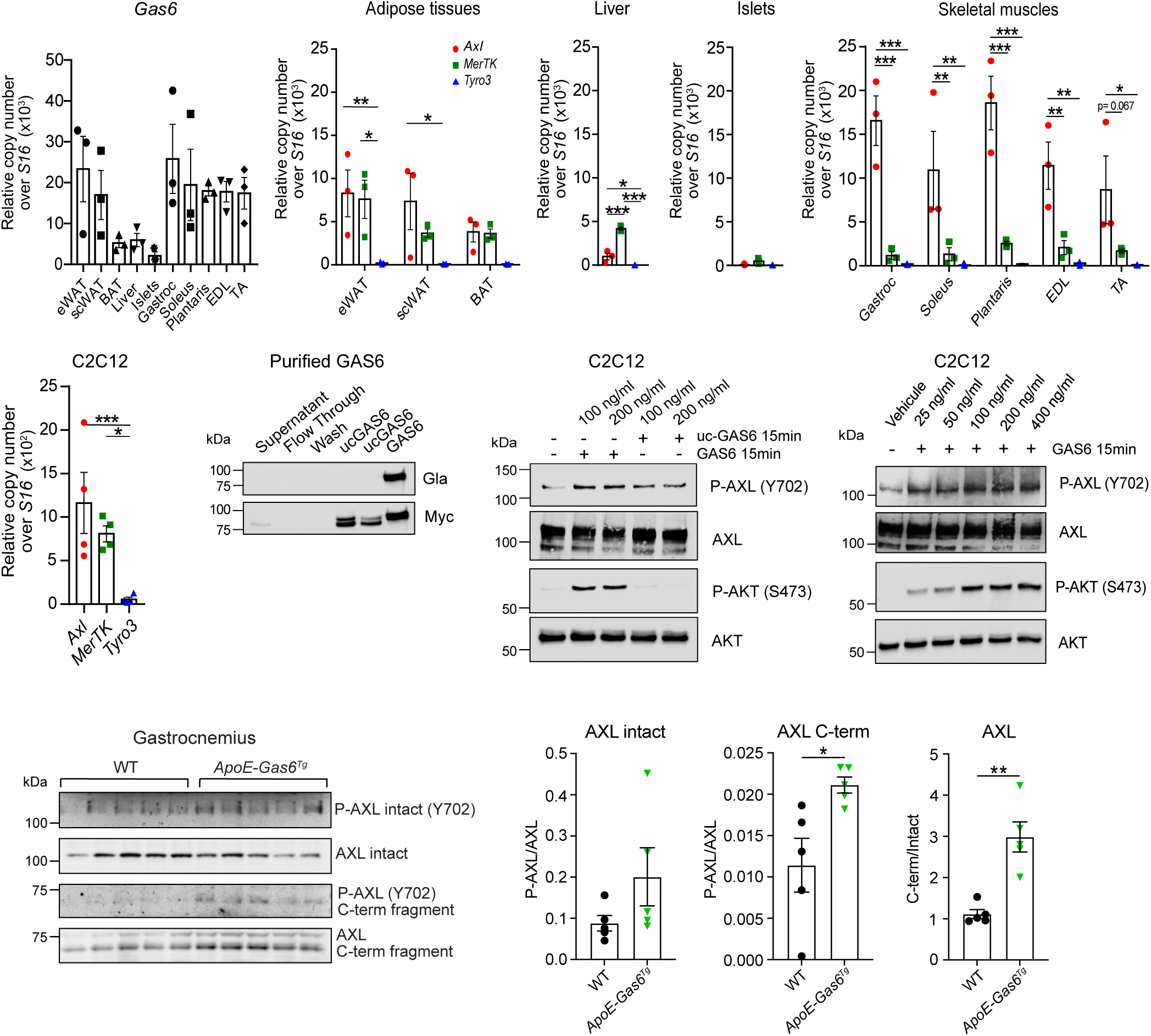
GAS6 activates AXL in muscle cells. **(A)** *Gas6* and **(B-E)** *Axl, Mertk* and *Tyro3* (TAM receptors) expression in WT (C57BL/6J) mouse tissues measured by quantitative PCR (qPCR) (n=3). eWAT: epididymal white adipose tissue; scWAT: subcutaneous white adipose tissue; BAT: brown adipose tissue; Gastroc: gastrocnemius; EDL: extensor digitorum longus; TA: tibialis anterior. Results are represented as copy number normalized to *S16.* **(F)** TAM receptors expression in C2C12 cells was measured by qPCR (n=4). Results are represented as copy number normalized to *S16.* **(G)** Recombinant GAS6 carboxylation and expression at different steps of GAS6 purification, tested using α-Gla antibodies by Western blot analysis. α- Myc was used to identify GAS6-Myc-6×His protein. **(H)** Phosphorylation of AXL (Y702) and AKT (S473) in C2C12 myotubes treated with uncarboxylated (uc-GAS6) or carboxylated GAS6 (GAS6: 100 and 200 ng/mL) for 15 minutes. Total AXL and total AKT were used as loading controls. **(I)** Dose-response of AXL and AKT phosphorylation in C2C12 cells treated with GAS6 for 15 minutes. **(J)** Phosphorylation of AXL (Y702) and total AXL levels in gastrocnemius muscle from WT and *ApoE-Gas6^Tg^* mice at 3 months of age (n=5). **(K)** Quantification of phosphorylated intact AXL over total intact AXL. **(L)** Quantification of phosphorylated cleaved AXL (C-terminal AXL fragment (C-term)) over total cleaved AXL. **(M)** Quantification of total cleaved AXL normalized over the amount of total intact AXL. Results represent mean ± SEM, \**P* < 0.05, \*\**P* < 0.01, \*\*\**P* < 0.001 by one-way ANOVA with Bonferroni’s post test **(B-F)** or Student’s *t* test **(K-M)**. Western blot are representative experiments of at least 3 independent experiments.

For this purpose, we selected the C2C12 myoblast cell line which can be differentiated into myotubes, respond robustly to insulin (31) and express *Axl* (**Fig.4F**). Since AXL is the TAM family member with the highest affinity for GAS6 (10), we focused our study on this receptor. Recombinant mouse GAS6 was produced and purified in HEK293 cells in the presence of vitamin K to induce its γ-carboxylation (GAS6) or warfarin to produce uncarboxylated GAS6 (uc-GAS6) (**Fig.4G**). Phosphorylation of AXL and its downstream target AKT increased when myotubes were treated with GAS6, but not with uc-GAS6 (**Fig.4H**), confirming that recombinant GAS6 was active, and that γ-carboxylation is necessary for its activity. In addition, GAS6 induces AXL and AKT phosphorylation in a dose-dependent manner with maximal activation between 100 and 200ng/mL (**Fig.4I**). Phosphorylation of AXL was increased in the muscle of *ApoE-Gas6^Tg^*mice and correlated with increased cleaved (i.e., C-terminal fragment) over intact AXL ratio, consistent with increased ligand-induced AXL shedding in the GAS6 overexpressing animals (**Fig.4J-M**). Together, these data show that GAS6 can activate AXL in muscle cells in vitro and in vivo.

### GAS6 suppresses insulin response in muscle cells

To determine if GAS6 and AXL could impact insulin signaling, C2C12 cells were pre-treated for 8 or 24 hours with GAS6 (200 ng/ml) or vehicle, followed by a 15-minute treatment with insulin. 24-hour long treatment with GAS6 significantly decreases by ∼2-fold IR phosphorylation induced by insulin (**Fig.5A-B**). An intermediate, but not significant reduction was observed with the 8-hour GAS6 treatment. The 8-hour GAS6 treatment also reduced AKT phosphorylation on S473 in response to insulin, but it reached significance only after cells were pre-treated with GAS6 for 24 hours (**Fig.5A-B**). Similar results were obtained in L6-GLUT4myc cells, a rat myoblastic cell line that stably expresses human GLUT4 tagged with a myc epitope (**Fig.S4A**). In this cellular model, 24 hours of GAS6 treatment also reduced insulin-induced translocation of GLUT4 to the plasma membrane (**Fig. S4B**).

**Figure 5.**
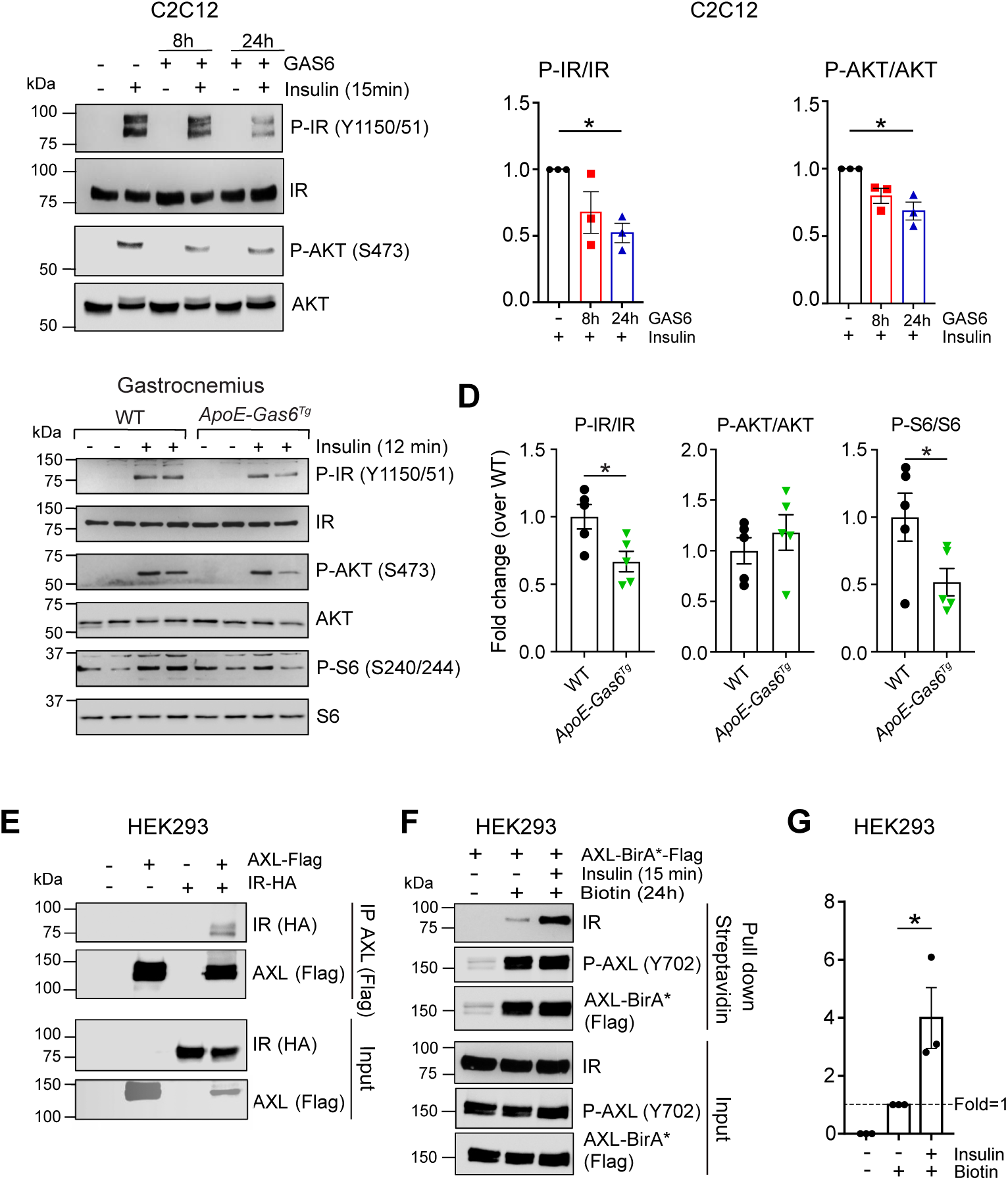
GAS6 suppresses insulin signaling in C2C12 and AXL interacts with the insulin receptor. **(A)** Phosphorylation of the insulin receptor (IR-Y1150/1151) and AKT (S473) assessed by Western blot in C2C12 cells treated with GAS6 (200 ng/ml) for 8h or 24h followed by insulin stimulation (100 nM) for 15 minutes. Total IR and total AKT were used as loading controls. **(B)** Quantification of phosphorylation levels of IR and AKT normalized over the amount of total protein (n=3). **(C)** Representative western blot showing the phosphorylation of the insulin receptor (IR; Y1150/1151), AKT (S473) and ribosomal protein S6 (S240/244) in gastrocnemius muscle of WT and *ApoE-Gas6^Tg^* mice 12 minutes after an i.v. injection of saline (-) or insulin (0.5U/kg). **(D)** Quantification of the phosphorylation levels normalized over the total amount of each protein (n=5). **(E)** Co-immunoprecipitation (IP) in HEK293 transfected with AXL-BirA*-Flag and IR-HA plasmids. **(F)** Proximity-dependent biotinylation (BioID) assay performed in HEK293 Flp-In T-REx cells expressing AXL-BirA*-Flag. **(G)** Quantification of endogenous IR signal detected by streptavidin pulldown and normalized over the biotin only condition. Results represent mean ± SEM. \**P* < 0.05, by one-way ANOVA followed by Bonferroni’s post test **(B, G),** or by unpaired, 2-tailed Student’s *t* test **(D)**. Western blot are representative experiments of 3 independent experiments.

Phosphorylation of IR and of ribosomal protein S6 (S240/S244), a downstream target of IR and AKT, was reduced in the muscle of *ApoE-Gas6^Tg^*mice compared to WT animals in response to an insulin injection (**Fig.5C-D**). No difference in the phosphorylation of AKT (S473) was observed, possibly due to difference in kinetic of activation of the pathway or in tissue insulin concentrations compared to the in vitro experiments. IR, AKT and S6 phosphorylation in response to insulin was normal in WAT and liver of *ApoE-Gas6^Tg^* mice (**Fig.S4C-F**). These results suggest that chronic exposure to GAS6 reduces the capacity of muscle cells to respond to insulin in vitro and in vivo.

### AXL interacts with the insulin receptor

It was previously reported that AXL can interact with other RTK, such as the EGFR, and rewire the signaling pathways elicited by these receptors (32). In HEK293 cells transfected with AXL-Flag and/or IR-HA expression vectors, IR was co-immunoprecipitated (co-IP) with AXL (**Fig.5E**). BioID, a proximity biotin labeling assay (33), was used to corroborate these results. HEK293 Flp-In T-REx cells expressing the AXL-BirA*-Flag fusion protein were treated with biotin for 24 hours, biotinylated AXL proximal proteins pulled-down with streptavidin Sepharose beads and analyzed by western blot. This approach suggests that IR forms a complex with AXL in mammalian cells (**Fig.5F**) and that the interaction is about 4-fold stronger when the cells are stimulated with insulin (**Fig.5G**). In these experiments performed in HEK293 cells and as previously reported (16), we observed that overexpressed AXL-BirA*-Flag was phosphorylated in the absence of ligand, indicating ligand-independent activation of AXL kinase activity. In C2C12, in which AXL activation depends on GAS6 (**Fig.4H-I**), insulin treatment did not impact AXL phosphorylation (**Fig.S4G**). Taken together, these results suggest that AXL and IR form a complex in an insulin-dependent manner.

### GAS6 reprograms insulin-dependent signaling pathways in muscle cells

To uncover signaling pathways modulated by GAS6 and impacting insulin signaling, we compared the skeletal muscle (gastrocnemius) transcriptome of *Gas6*^+/+^ and *Gas6*^-/-^ mice. This analysis revealed that only 17 genes were significantly dysregulated in the absence of GAS6 (*P-adj* < 0.05; **Fig.S5A**). The mRNA encoding for IR (*Insr*) was reduced by ∼30% in the muscle of *Gas6*^-/-^ mice, however, IR protein level was not affected in the same tissue and in WAT and liver (**Fig.S5B-D**). These results show that GAS6 has a limited effect on muscle gene expression, suggesting that it may influence insulin signaling through post-translational mechanisms.

Quantitative phosphoproteomics was next used to decipher the molecular impact of GAS6 on insulin-dependent signaling. C2C12 cells were treated with GAS6 or vehicle for 24 hours, before being stimulated or not with insulin for 15 minutes. Phosphoserine and phosphothreonine containing peptides were enriched using TiO_2_ chromatography, labeled with Tandem Mass Tags (TMT) and combined to be analyzed by high-resolution liquid chromatography–tandem MS (LC-MS/MS) to quantify the relative abundance of phosphopeptides modulated by GAS6, insulin or their combination (**Fig.6A**) (34). The experiment was independently repeated three times, median normalized, and batch effects were corrected. Only the peptides detected and quantified in all the experiments were considered for further analysis. Using this approach, quantitative data were obtained for a total of 1491 unique phosphopeptides, originating from 712 different proteins (**Supplementary Table 3)**. We then performed a series of pairwise comparisons and only included peptides that were modulated by at least 1.5-fold between two given conditions with a *P* value ≤ 0.05 (**Fig.S6A** and **Supplementary Table 4**).

**Figure 6.**
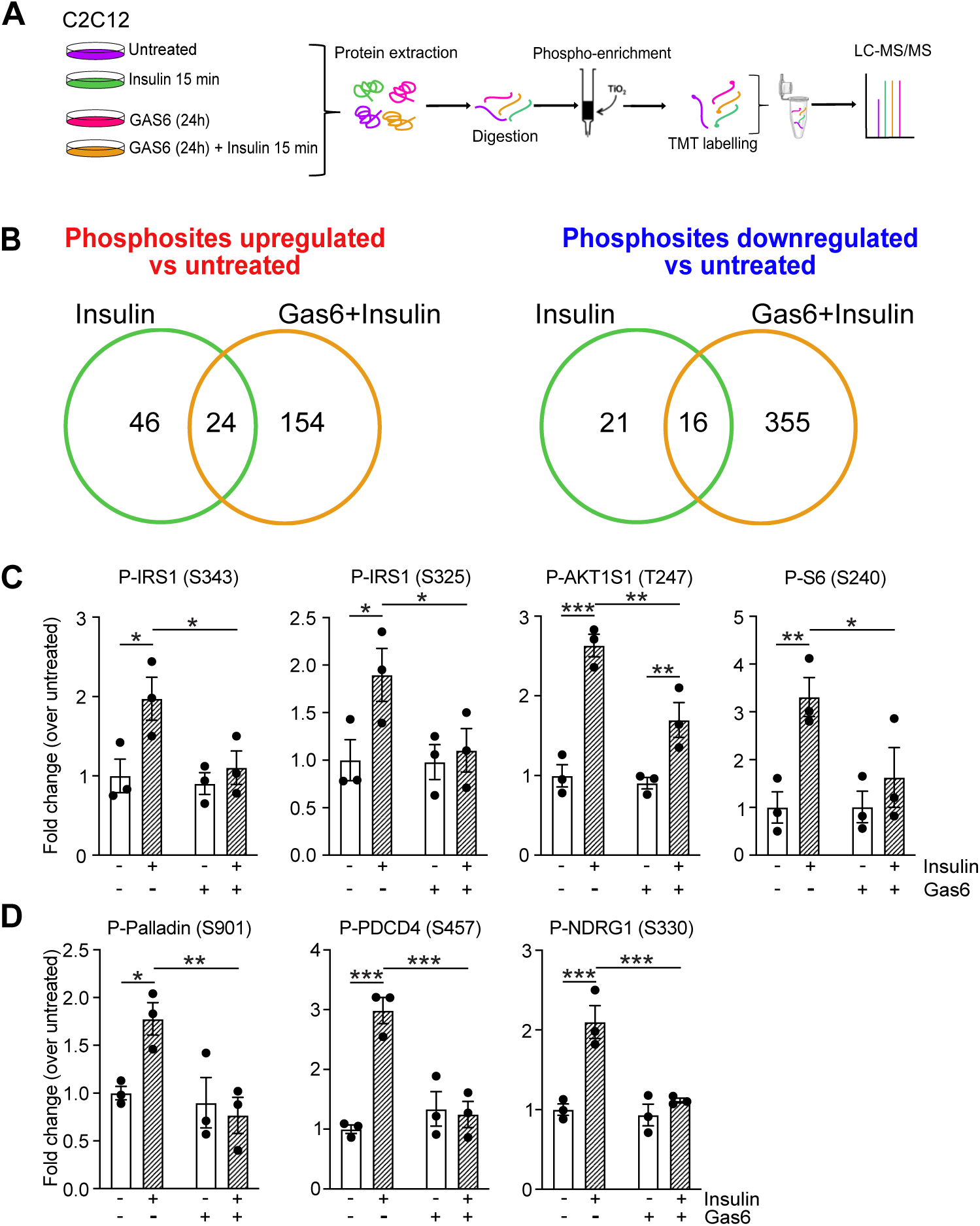
GAS6 reprograms the insulin-dependent phosphoproteome in muscle cells. **(A)** Schematic representation of the experimental design, including phospho-enrichment, TMT labelling and LC-MS/MS analysis of C2C12 treated for 24 hours with GAS6 (200ng/ml) followed by 15 minutes of insulin stimulation (100nM). The phosphosproteomic screen was performed on samples from 3 independent experiments. **(B)** Venn diagram of phosphosites upregulated and downregulated by insulin or GAS6 + insulin treatments in comparison to untreated condition (i.e, no insulin). **(C)** Phosphorylation levels of insulin and AKT downstream targets as measured in the phosphoproteomic analyses (n=3). Results represent mean ± SEM. \**P* < 0.05, \*\**P* < 0.01, \*\*\**P* < 0.001 by 2-way ANOVA followed by Fisher’s LSD post test.

Only 54 phosphopeptides were found to be significantly modulated by the 24-hour GAS6 pre-treatment compared to untreated cells (**Fig.S6A**). A larger number of phosphorylation sites were up- or down-regulated following the 15 minutes insulin treatment, both in the presence and absence of GAS6 pretreatment (**Fig.6B and Fig.S6A**). Interestingly, the total number of unique phosphopeptides changed by insulin following GAS6 pretreatment was approximately 6-times higher when compared to insulin alone (549 vs. 107 phosphopeptides; **Fig.6B and Fig.S6A**). However, the overlap between these two groups was limited, with only 24 phosphopeptides upregulated and 16 downregulated by insulin both in the presence and absence of GAS6 (**Fig.6B**), suggesting that GAS6 treatment significantly modifies the insulin-dependent phosphoproteome in C2C12 cells.

A total of 70 phosphosites, contained in 49 different proteins, were upregulated by insulin alone (**Fig.6B; Fig.S6A and Supplementary Table 5)**. Several phosphosites on proteins associated with insulin signaling and the AKT pathway, such as IRS1 (S343 and S325), AKT1S1/PRAS40 (T247) and ribosomal protein S6 (S240) were increased by insulin alone, but reduced in cell pretreated with GAS6 (**Fig.6C**). In addition, other known AKT substrates, including palladin (S901), PDCD4 (S457) and NDRG1 (S330), were also negatively affected by GAS6, confirming that GAS6 inhibits AKT signaling in these cells (**Fig.6D**).

We next concentrated our analysis on the phosphorylation events differentially affected when comparing insulin with insulin in combination with GAS6 (*P*-adj <0.05; fold-change ≥ 1.5; **Supplementary Table 6)**. Reactome pathways enrichment analysis showed that proteins whose phosphorylation was decreased in response to insulin in the presence of GAS6, were mainly associated with RNA metabolism, RhoGTPase signaling, and signal transduction by growth factor receptors (**Fig.S6B-C**).

The group of proteins containing phosphorylation sites that are increased by insulin in the presence of GAS6 displayed a very strong enrichment in pathways involving small GTPases of the Rho, Rab, RAC and CDC42 families (**Fig.S6D-E**). Several proteins involved in Rho GTPases signaling contained multiple phosphorylation sites that were up- and down-regulated by insulin in the presence of GAS6 (**Fig.S6C and S6E**). While there were no specific changes in phosphorylation of RAC1 and CDC42 GTPases, GAS6 altered the phosphorylation of their GEFs, in response to insulin, respectively DOCK7 and ARHGEF40, as well as their effectors, the kinase of the PAK family (PAK1, PAK2 and PAK4) (**Fig.S6D-E**).

The group of proteins containing phosphorylation sites upregulated by insulin in the presence of GAS6 was also enriched in pathways involved in membrane trafficking and vesicle-mediated transport (**Fig.S6D-E**). These included SEC31A (S526), a component of the COPII protein complex responsible for vesicle budding from the endoplasmic reticulum (ER) and adaptor-associated protein kinase 1 (AAK1; S635) involved in clathrin-mediated endocytosis. Several proteins in this group have a function in endocytosis and endosomal trafficking which concur with the enrichment of Rab family small GTPases pathways known to control vesicular membrane trafficking to endosomes (**Fig.S6E)**. Altogether, this phosphoproteomic study demonstrates that GAS6 significantly modifies the molecular response to insulin.

### AXL signaling increases insulin receptor endocytosis in response to insulin

Analysis of the phosphoproteome uncovered an endocytosis and endosomal trafficking pathway specifically modulated by insulin in the presence of GAS6 and implicating the Rab GTPases, their effectors, and other regulators of endocytosis (**Fig.7A-B and Fig.S7A**). Endocytosis regulates signaling from RTKs, including IR (35) and Rab GTPases are known to play a critical role in endosomal trafficking (36). Formation of late endosomes consists of a transition of Rab5 to Rab7a at their surface. The quantitative proteomic data indicate that Rab7a (S72) itself and TBC1D15 (S205), one of its GAPs, are more phosphorylated following insulin stimulation in the presence of GAS6 (**Fig.7B and Fig. S7A**). Phosphorylation of Rab7a on S72 was previously shown to facilitate the dynein-driven transport of EGFR, another RTK, toward the perinuclear region (37). These observations prompted us to investigate whether GAS6- and AXL-elicited signaling could impact insulin receptor (IR) endocytosis and intracellular trafficking.

**Figure 7.**
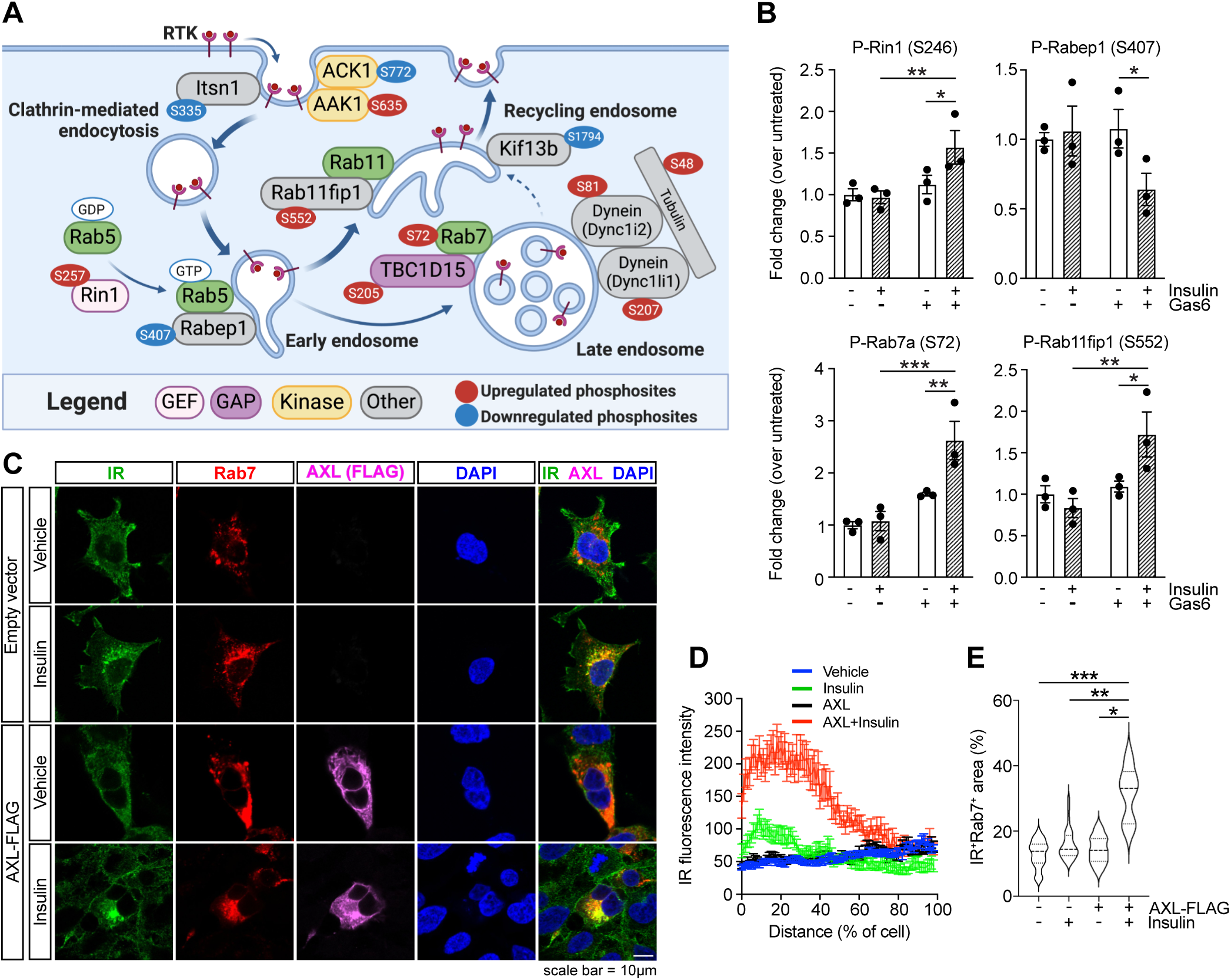
AXL increases insulin receptor endocytosis in response to insulin. **(A)** Schematic of the endocytosis pathway involving Rab proteins, their regulators, and their effectors, specifically affected by insulin and GAS6 co-treatment. Upregulated and downregulated phosphosites by insulin in combination with GAS6 are represented in red and blue respectively (Figure created using BioRender). **(B)** Phosphorylation levels of proteins highlighted in **(A)** as measured in the quantitative phosphoproteomics analysis (n=3). **(C)** Representative confocal microscopy images of HEK293 cells transfected with Rab7-mScarlet and AXL-Flag or empty vector, and treated or not with 100 nM insulin for 15 minutes. Cells were stained for AXL-Flag (Alexa Fluor 633/magenta) and endogenous insulin receptor (IR; Alexa-Fluor 488/green). DAPI was used to stain nuclei. Scale bar, 10 µm. **(D)** Quantification of IR fluorescence signal from the nucleus to the plasma membrane normalized to cell size (n=30 cells per conditions, from 3 independent experiments). **(E)** IR and Rab7 colocalization fluorescence signal normalized over cell area. Results represent mean ± SEM **(B, D)**, or median with quartiles **(E)**. \**P* < 0.05, \*\**P* < 0.01, \*\*\**P* < 0.001 by 2-way ANOVA followed by Fisher’s LSD post test **(B)**, or by one-way ANOVA with Bonferroni’s post test **(E)**.

Visualizing IR trafficking in myoblasts or myotubes is challenging due to cell shape, size and autofluorescence. We therefore used HEK293 cells, which express endogenous IR and respond to insulin (38) but do not express AXL (39). These cells were transfected with mScarlet-tagged Rab7a to visualize late endosomes, and with AXL-Flag or an empty vector, treated with insulin for 15 minutes and the cellular localization of endogenous IR and Rab7a was assessed by immunofluorescence (**Fig.7C** and **Fig.S7B**). Stimulation with exogenous GAS6 was not required since AXL is already robustly phosphorylated in the absence of ligand in this experimental model (see **Fig.5F** and (16)). In the absence of AXL and insulin stimulation, IR was detected mostly at the level of the plasma membrane (**Fig.7C-D**). Following a 15-minute treatment with insulin, IR is partly redistributed from the membrane toward the perinuclear region in part in Rab7a-positive endosomes (**Fig. 7C-E**). In the presence of AXL and in the absence of insulin, the IR exhibits a cellular distribution similar to the one observed in untransfected cells. Strikingly, in the presence of AXL and following insulin stimulation, the intracellular accumulation of IR in the Rab7a-late endosomes was significantly increased (**Fig.7C-E**).

The impact of GAS6 and insulin on IR trafficking was next assessed in vivo by immunofluorescence staining on gastrocnemius sections. In wildtype mice injected with saline, the IR appears to be localized at the membrane of muscle myofibers, but following an insulin injection, the membrane localized IR was significantly reduced (**Fig.8A-D**). In *ApoE-Gas6^Tg^*muscle, membrane-localized IR is further reduced following insulin injection (**Fig.8A-B**). In contrast, in *Gas6*^-/-^ mice injected with insulin, the membrane-associated IR remain higher compared to wildtype mice in the same conditions (**Fig.8C-D**). Altogether, these data suggest that AXL and GAS6 signaling potentiates insulin-induced IR endocytosis.

**Figure 8.**
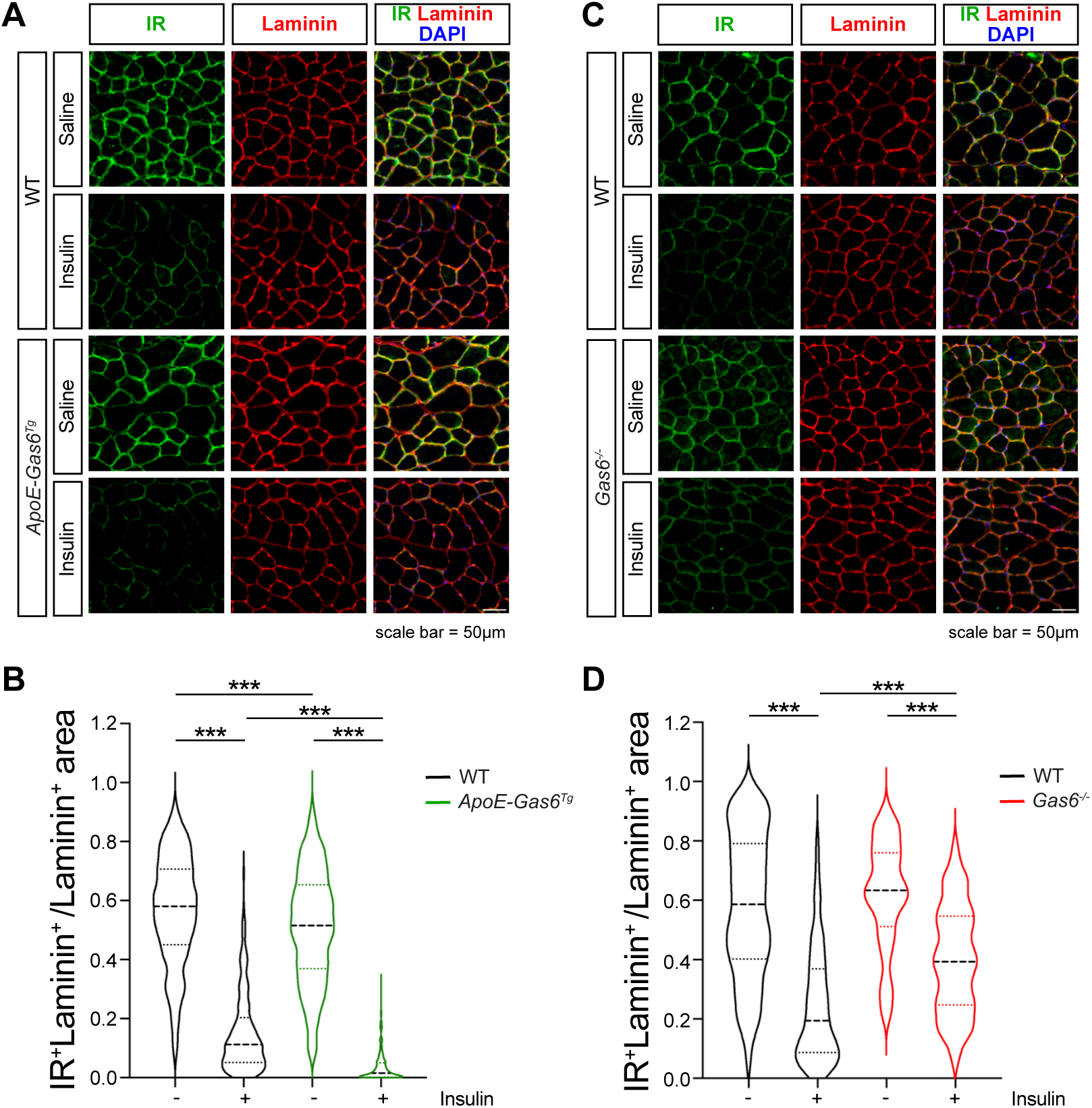
GAS6 regulates insulin receptor membrane localization in muscle in vivo. **(A)** Representative confocal microscopy images of gastrocnemius muscle sections from WT and *ApoE-Gas6^Tg^*mice 12 minutes after an i.v. injection of saline (-) or insulin (0.5U/kg), stained for insulin receptor (IR; Alexa-Fluor 488/green) and Laminin (DyLight-650/red). DAPI was used to stain nuclei. Scale bar, 50 µm. **(B)** Quantification of the colocalization signal of IR and Laminin at cell surface in individual myofiber (100 to 200 myofibers from 2-5 mice were analyzed per conditions). **(C)** Representative confocal microscopy images of gastrocnemius muscle sections from WT and *Gas6*^-/-^ mice 12 minutes after an i.v. injection of saline (-) or insulin (0.5U/kg), stained as in **(A)**. Scale bar, 50 µm. **(D)** Quantification of the colocalization signal of IR and Laminin at cell surface in individual myofiber (100 to 200 myofibers from 4-5 mice were analyzed per conditions). Results represent median with quartiles. \*\*\**P* < 0.001 by one-way ANOVA with Bonferroni’s post test.

## DISCUSSION

A few studies have investigated the role of GAS6 and AXL in obesity and glucose metabolism using mouse models, leading to conflicting results. For instance, *Gas6*^-/-^ mice on a mixed genetic background (Swiss:129SV, 50:50) accumulate less WAT compared to *Gas6*^+/+^ mice when fed a high-fat diet (40). Similarly, treatment with R428 (25mg/kg twice a day), a pharmacological inhibitor of AXL, resulted in significantly reduced weight gain and subcutaneous and gonadal fat mass in C57Bl/6 mice fed a HFD (41). In contrast, *Axl*^-/-^ mice on a different mixed genetic background (C57Bl/6:129S1, 50:50) fed a HFD displayed no difference in adipose depot mass compared to controls (42). Importantly, glucose tolerance and insulin sensitivity were not investigated in detail in any of these studies. R428 is a pharmacological inhibitor of AXL which can inhibit other tyrosine kinase receptor at high doses, including IR, EGFR, MerTK and TYRO3. Off-target effects may therefore potentially explain the reported impact of R428 treatment on fat depots in mice.

Here, we extensively characterize the glucose and metabolic phenotype of *Gas6*^-/-^ mice on two inbred genetic backgrounds. Our data show that in these well-controlled conditions, *Gas6*-deficiency does not impact food intake, energy expenditure, body weight and adipose tissue mass on normal chow diet or HFHS diet. However, genetic ablation of GAS6 was associated with reduced blood glucose and improved glucose tolerance and insulin sensitivity. To precisely dissect the impact of GAS6-signaling on insulin-target tissues, it will be interesting to generate muscle, WAT, and liver specific knockout of *<u>Gas6</u>* <u>and</u> *Axl* in mice.

Using two different approaches, co-IP and BioID, we provide evidence that AXL and IR form a complex in mammalian cells. AXL has been shown to heterodimerize and/or to crosstalk with other RTK including HER2, EGFR, MET, VEGFR-2 and PDGFRβ (32; 43-45). In these instances, AXL appears to potentiate or diversify the signaling pathways emanating from its partner RTK. The IR-AXL crosstalk we describe here is different since it results in a downregulation of classical IR signaling pathways and an increased trafficking of IR to the endosomes. Moreover, the IR and AXL interaction appears to be positively regulated by insulin, the IR ligand, suggesting that AXL preferentially associate with the IR in its active conformation.

Our phosphoproteomic data show that chronic treatment of muscle cells with GAS6 significantly affected the phosphorylation of several proteins involved in vesicular trafficking in response to insulin, including proteins of Rab GTPases (e.g., Rab7a S72). The phosphorylation of multiple kinases appears to be specifically modulated by insulin in the cell pretreated with GAS6 for 24h. Some of these kinases have previously been implicated in the regulation of endocytosis (e.g., Aak1 and Ack1) or in RHO GTPase regulation (e.g., Rock2, Pak1, 2 and 4, Peak1, Pnk1 and Pnk2). It is plausible that the activation or inactivation of this specific set of kinases explains the differential phosphorylation of substrates induced by insulin in presence of GAS6.

Other results presented here revealed that AXL increases the internalization of the IR and its localization in Rab7a-positive endosomes after insulin stimulation. This change in localization corroborate studies showing a decrease in the levels of IR at the plasma membrane in obese and diabetic conditions (46; 47), although downregulation of the IR is not always present in pathological insulin resistance (48). Interestingly, a subclass of IR mutations causing severe insulin resistance and T2D in humans is associated with an abnormal enrichment of the IR in the Rab7a-positive endosomes (49), suggesting that premature or excessive IR endocytosis can contribute to insulin resistance in humans. In addition, RTK signaling can still occur during endocytosis, since their trafficking to endosomes disrupts or facilitates their proximity to downstream signaling effectors (50). Finally, it is also possible that GAS6/AXL signaling slows down the kinetic of IR recycling from endosome to the membrane thereby reducing insulin response.

In conclusion, our results establish a new role for GAS6 and AXL in the remodeling of insulin signaling in muscle cells and in the development of insulin resistance. This study paves the way for the investigation and development of potential therapeutic approaches using existing pharmacological inhibitors of GAS6/AXL signaling to improve insulin sensitivity.

## ACKNOWLEDGEMENTS

We thank Drs Mark Blostein for providing the *Gas6*^-/-^ mice on C57Bl/6J background and Dr Amélie Robert for generating the pmScarlet-C1-Rab7a plasmid. We also thank the staff of the IRCM Bioinformatics, Proteomics, Microscopy, Molecular Biology and Histology Core Facilities for their technical support.

## AUTHOR CONTRIBUTIONS

MF and JL conceived, designed the project, and supervised CS and AG. CS, AG, JL, MP, DF, JB and MF collected and analyzed data. JL and CS assembled the figures. PC provided the *Gas6*-/- mice. JFC provided unique reagents and critical advice for experiments. CS, JL, and MF wrote the manuscript with suggestion from the other authors.

## Guarantor Data Access and Responsibility Statement

Dr. Mathieu Ferron is the guarantor of this work, had full access to all the data, and takes full responsibility for the integrity of data and the accuracy of data analysis.

