## Supplemental Material and Figures for "GAS6 and AXL promote insulin resistance by rewiring insulin signaling and increasing insulin receptor trafficking to endosomes"

### SUPPLEMENTAL RESEARCH DESIGN AND METHODS

#### Mice

*Gas6*$^{-/-}$ mice on a C57BL/6J genetic background were generated in Dr Peter Carmeliet laboratory [1]. *Gas6*$^{-/-}$ mice on a FVB/N genetic background were generated by Dr Jean-François Côté laboratory [2]. *ApoE-Gas6*$^{Tg}$ transgenic mice were generated in our laboratory. The sequence of mouse *Gas6-Myc-His* cDNA was cloned into pLiv.7 plasmid containing liver-specific human APOE promoter and hepatic enhancers [3] (**Supplemental Table 1**). After excision from pLiv.7 plasmid with KpnI and XhoI restriction enzymes *the ApoE-Gas6* transgenic construct was purified and microinjected into C57BL6/J mouse fertilized oocytes. *ApoE-Gas6*$^{Tg}$ were maintained on a pure C57BL6/J genetic background. Transgenic mice were identified by genotyping using specific primers (**Supplemental Table 1**). All strains were maintained in an IRCM specific pathogen-free animal facility under 12-hour dark/12-hour light cycles. Mice were fed ad libitum a normal chow diet (Teklad global 18% protein rodent diet; 2918; Envigo), unless otherwise specified. Male mice were used in all experiments.

#### Cell Lines

C2C12 cells (ATCC) were cultured as myoblasts (undifferentiated form) in DMEM, supplemented with 10% fetal bovine serum (FBS) and penicillin/streptomycin (P/S). Differentiation of C2C12 myoblasts into myotubes was induced by replacing media, after cells reached approximately 70-80% of confluency, with DMEM containing 2% horse serum for 5 days and changing media every 2 days. HEK293 cells (ATCC) were cultured in EMEM with 10% heat inactivated FBS and P/S. HEK293 Flp-In T-REx cells (Thermo Fisher) expressing or not AXL-BirA*-Flag were obtained from Dr Jean-François Côté, and maintained in DMEM, 10% FBS and P/S. L6-GLUT4myc rat myoblast cells (Kerafast) were cultured as myoblasts in αMEM supplemented with 10% FBS and P/S with 5µg/ml blasticidin. All the cells were cultured at 37°C, with 5% $CO_2$.

#### DNA Constructs

pcDNA5-AXL-BirA*-Flag was generated by inserting the human AXL cDNA upstream of BirA*-FLAG in the pDEST-pcDNA5-BirA-FLAG-Nterm, obtained from Dr. A.-C. Gingras. Mouse GAS6-Myc-6xHis was generated by PCR amplification of mouse *Gas6* and cloning in the EcoRI and XbaI sites of pcDNA5/FRT/TO-pDEST (Thermo Fisher). pcDNA3-IR-HA was generated by PCR amplification of the insulin receptor (IR) and cloning with HA-tag in the HindIII and EcoRI restriction sites of pcDNA3 plasmid. pmScarlet-C1-Rab7a was generated by PCR amplification of Rab7a from mRFP-C3-Rab7a (Addgene; 14436) and cloning in the HindIII and ApaI sites of pmScarlet-C1 plasmid. Sequences of the primers used for cloning are included in **Supplemental Table 1**.

**Production and Purification of Recombinant GAS6**

HEK293 Flp-In T-REx cells were transfected using Lipofectamine 2000 (Invitrogen) with pcDNA5-pDEST-mGAS6-Myc-6xHis and pOG44 (Thermo Fisher) which expressed the Flp recombinase, to generate a cell line expressing recombinant mouse GAS6 in a Tetracycline-inducible manner. Cells were then selected with hygromycin for 12 days. After expansion, cells were treated for 48 hours with tetracycline (1µg/ml) and vitamin $K_1$ (22µM, Millipore Sigma) in DMEM media without serum. The supernatant was collected, centrifuged and filtered to eliminate cells and debris. The recombinant GAS6 protein was purified via His-tag by affinity chromatography using a 1mL HisTrap HP column (Cytiva) connected to an AKTAprime FPLC system (GE). The supernatant was loaded onto the HisTrap HP column pre-equilibrated with binding buffer (20 mM sodium phosphate, 0.5 M NaCl, 20 mM imidazole, pH 7.4) and washed with 10 column volumes of binding buffer at a flow rate of 1 mL/min. Elution was performed using buffer containing: 20 mM sodium phosphate, 0.5 M NaCl and 500 mM imidazole (pH 7.4). Purified proteins were dialyzed in PBS 1X and quantified using mouse GAS6 ELISA (DuoSet ELISA, R&D Systems, DY986) following manufacturer's instructions. Uncarboxylated GAS6 was produced following the same procedure except that warfarin (50µM) instead of vitamin $K_1$ was added to the media during the induction step.

### Co-Immunoprecipitation and Western Blot

Proteins were extracted with lysis buffer containing 20mM Tris-HCl (pH 7.5), 150mM NaCl, 1mM EDTA (pH 8.0), 1mM EGTA, 2.5mM NaPyrophosphate, 1mM β-glycerophosphate, 10mM NaF, 1% Triton, 1 mM vanadate, 1mM phenylmethylsulfonyl fluoride (PMSF) and protease inhibitors (4693132001; Roche Diagnostics) as previously described [4] and quantified using Bradford assay (Biorad). For co-immunoprecipitation, HEK293 cells were transfected with pcDNA5-AXL-BirA*-Flag and pcDNA3-IR-HA using Lipofectamine 2000 (Invitrogen). After two days, cells were harvested, and 500 µg of protein lysate was incubated with anti-Flag agarose beads (Sigma) for 2 hours at 4°C with rotation, followed by 4 washes with lysis buffer. Laemmli buffer was added to co-immunoprecipitated proteins or input samples which were then heated for 10 minutes at 70°C. Proteins were detected by Western blot with the indicated primary antibodies, overnight at 4°C and quantified by densitometry analyses using Image Lab software (version 5.0; Bio-Rad Laboratories). Antibodies used in this study are listed in **Supplemental Table 2**.

### Streptavidin Pull-Down

HEK293 Flp-In T-REx cells expressing AXL-BirA*-Flag in a tetracycline-inducible manner were cultured during 24 hours with biotin (50µM) and tetracycline (1µg/ml). A negative control without biotin was also included. Cells were cultured in 0.5% FBS DMEM overnight, then incubated in serum-free medium (with 0.1% BSA and 10 mM HEPES) for 2 hours followed by 15-minute stimulation with insulin (100 nM). Proteins were extracted with the lysis buffer described above and 2mg of protein lysate was incubated with anti-streptavidin beads (Sigma) for 3 hours at 4°C with rotation. Beads were then washed 6 times with lysis buffer. Biotinylated proteins or input samples were diluted in Laemmli buffer supplemented with 20mM DTT and 2mM biotin. Samples were boiled for 15 minutes at 95ºC and used for western blot analysis.

### Immunofluorescence Confocal Microscopy

HEK293 cells were transfected with pcDNA5-AXL-BirA*-Flag and/or pmScarlet-C1-Rab7 and replated on poly-L-Lysine coated glass coverslips the next day. Two days later, cells were fixed with 4% paraformaldehyde (PFA) for 15 minutes. Unspecific binding was blocked with 5% Normal Donkey Serum in PBS containing 0.3% Triton for 1h at room temperature. Fixed cells were then incubated with rabbit anti-insulin receptor β and mouse anti-FLAG antibodies diluted in PBS, 1% BSA and 0.1% Triton, overnight at 4°C (**Supplemental Table 2)**. Cells were then incubated with Alexa Fluor 488-conjugated donkey anti-rabbit and Cy3-conjugated donkey anti-mouse secondary antibodies for 1h at room temperature (**Supplemental Table 2**). DAPI was used to stain nuclei. Images were acquired with Zeiss LSM700 confocal microscope with a 63X oil objective. We used Image J software to quantify Insulin Receptor (IR) and Rab7 localization. For each different condition, IR (Alexa488) and Rab7 (Scarlet) signal intensities were measured across the cell from the nucleus to the plasma membrane and normalized to cell size (n=30 cells per condition, from 3 independent experiments). Colocalization percentage was measured as IR and Rab7 overlapping area for each cell over cell area.

For in vivo imaging of IR localization, gastrocnemius muscles were dissected and directly embedded with OCT compound. Ten µm sections were fixed in 4% paraformaldehyde for 10 min, permeabilized for 10 min in 1% normal donkey serum in PBS containing 0.4% Triton and further blocked in 5% normal donkey serum in PBS containing 0.1% Triton for 1h at room temperature. Sections were incubated with rabbit anti-insulin receptor β primary antibody overnight at 4°C diluted in PBS, 1% normal donkey serum and 0.1% Triton (Supplemental Table 2). Sections were incubated with Alexa Fluor 488-conjugated donkey anti-rabbit secondary antibody and subsequently with Laminin DyLight 650 for 1h at room temperature (Supplemental Table 2). DAPI was used to stain nuclei. Images were acquired with Zeiss LSM710 confocal microscope with a 20X objective. Image J software was used to quantify the colocalization area of Insulin Receptor (IR) and Laminin for each myofiber (100 to 200 myofibers were analyzed per condition from 2-5 mice).

**Gene Expression**

Tissue and cell total RNA extraction and isolation were performed as previously described [5]. Total mRNA was treated with DNaseI (Invitrogen) and reverse transcribed with random hexamers and oligo dT primers using M-MLV reverse transcriptase (Invitrogen). Quantitative real-time PCR was performed using PowerUp SYBR Green Master Mix (A25741; Applied Biosystems) with gene-specific primers (**Supplemental Table 1**) on a ViiA7 Real-Time PCR System (Applied Biosystems). *Gas6* and TAM receptors gene copy numbers were calculated with a mouse genomic DNA standard curve and variation between biological replicate was normalized to *S16* (*Rps16*) expression.

**F4/80 Immunohistochemistry**

Epididymal adipose tissues were fixed in 10% formalin overnight at 4°C. Five µm paraffin sections were rehydrated and subjected to antigen retrieval with 10 mM citrate buffer, pH 6.0 at 95°C for 10 minutes. Slides were then stained with rabbit anti-mouse F4/80 antibody (**Supplemental Table 2**) overnight at 4°C in a humid chamber. Detection was performed with anti-rabbit Vectastain ABC-HRP Kit (Vector Laboratories, #PK-4001) and NovaRED substrate Kit (Vector Laboratories, #SK-4800). Sections were counterstained with hematoxylin, dehydrated and mounted with DPX Mount. For each mouse, 5 sections of epididymal adipose tissue were analyzed, and 6-11 mice were analyzed per genotype and diet. Images were acquired using 20X objective, Leica DM4000B microscope and the OsteoMeasure Analysis System (Osteometrics).

**Quantification of GLUT4myc Translocation**

Optical detection of GLUT4 translocation at the cell surface was performed using L6 myoblast cell line expressing stably a rat GLUT4 with a human myc-epitope (L6-GLUT4*myc*; Kerafast) as previously reported [6].

**RNA-Sequencing**

RNA from gastrocnemius of *Gas6*$^{+/+}$ and *Gas6*$^{-/-}$ mice were extracted using TRIzol reagent (Thermo Fisher). Elimination of DNA and further RNA purification were performed using Zymo-Spin IIICG columns according to the manufacturer's instructions (ZymoResearch, Irvin, CA). RNA libraries were prepared from 1000ng of total RNA.

Poly(A)+ transcripts were enriched using the NEBNext® Poly(A) mRNA Magnetic Isolation Module (New England Biolabs) and libraries were prepared with RNA Hyperprep Kit (KAPA). Libraries size distribution was assessed on a 2100 bioanalyzer (Agilent Technologies) and quantified by qPCR. Equimolar libraries were sequenced in paired-end reads (PE50) on a Novaseq 6000 system (Illumina) with a SP flowcell and an average coverage of 50M fragments per library. The quality of the raw reads was assessed with FASTQC v0.11.8. After examining the quality of the raw reads, no trimming was deemed necessary. The reads were aligned to the GRCm38 genome with STAR v2.7.6a with more than 82% of reads uniquely mapped. Raw counts were computed using FeatureCounts v1.6.0 based on Ensembl mouse reference genome v101. Differential expression was performed using DESeq2 R package and 18 differentially expressed genes (DEGs) were obtained using p-adjusted <0.05. DEGs heatmap was drawn based on z-score using Morpheus (Broad Institute).

**Protein Digestion for Phosphoproteome**

C2C12 cells were incubated in serum-free medium with or without recombinant GAS6 (200ng/ml) for 24 hours. Samples were treated with insulin (100 nM) for 15 minutes or a vehicle and collected in lysis buffer. A total of 2 mg of protein lysate per condition were reduced with 9 mM dithiothreitol at 37ºC for 30 minutes and, after cooling for 5 minutes, alkylated with 17 mM iodoacetamide at room temperature for 30 minutes in the dark. Nucleic acids were digested for 2 hours at 37ºC with benzonase in the presence of 2mM $MgCl_2$. The protein digestion was performed on-bead using the protein aggregation capture method to clean the samples from detergents and reagents. Proteins bound to SeraMag beads (Cytiva) were digested with Lys-C/Trypsin (Promega), in 50mM triethylammonium bicarbonate (TEAB) overnight at 37ºC.

**Isobaric Peptide Labeling and LC-MS/MS**

Protein digests were acidified with trifluoroacetic acid (TFA) and cleaned from reagents with an Oasis HLB extraction plate (Waters UK) following the manufacturer's instructions. The recovered peptides were lyophilized and subjected to phosphopeptides enrichment using the MagReSyn® $TiO_2$ beads (ReSyn Biosciences). All steps of the enrichment were performed according to the manufacturer's instructions. The enriched phosphopeptides were desalted with C18 ZipTip pipette tips (Millipore). Eluates were dried down in a vacuum centrifuge and reconstituted through agitation for 15 minutes in 25 µL of 100 mM Triethylammonium bicarbonate (TEAB). Isobaric labeling of the protein digests was performed using the 6-plex tandem mass tag (TMT) reagents following the manufacturer's instructions (Thermo Fisher Scientific). TMT labeled samples were pooled, acidified with TFA, cleaned with HLB (Waters Oasis HLB 96-well Elution Plate) following the manufacturer's instructions. Samples were evaporated to dryness and resuspended by agitation for 15 minutes in 20 µL of 1% ACN-1% FA. Then, 5 µL was loaded into a 75 µm i.d. × 150 mm Self-Pack C18 column installed in the Easy-nLC II system (Proxeon Biosystems). The buffers used for chromatography were 0.2% formic acid in water (buffer A) and 0.2% formic acid in acetonitrile (buffer B). Peptides were eluted with a three-slope gradient at a flowrate of 250 nL/min. Solvent B first increased from 2 to 27% in 140 min, then from 27 to 36% B in 40 minutes and finally from 36 to 89% B in 10 minutes. The HPLC system was coupled to Orbitrap Fusion mass spectrometer (Thermo Fisher Scientific) through a Nanospray Flex Ion Source. Nanospray and S-lens voltages were set to 1.3-1.8 kV and 60 V, respectively. Capillary temperature was set to 250 °C. Full scan MS survey spectra (m/z 360-1500) in profile mode were acquired in the Orbitrap at a resolution of 120,000 with a target value at 4e5. The 20 most intense peptide ions were fragmented in the HCD collision cell and analyzed in the Orbitrap at a resolution of 60,000 with a target value at 5e4 and a normalized collision energy at 38. A MS3 scanning was performed in the Orbitrap at a resolution of 30,000 upon detection of a neutral loss of phosphoric acid (48.99, 32.66 or 24.5 Th) in MS2 scans. Target ions selected for fragmentation were dynamically excluded for 10 sec.

**MS Data Analyses**

Raw data of $TiO_2$ phospho-enriched samples acquired by both neutral lost approach (NL1 and NL2) was analyzed with MaxQuant (version 2.0.1.0) by searching against Uniprot's mouse proteome database of sequences (August 2021 release) supplemented with MaxQuant's contaminants and reverse decoy sequences. All searches were performed using a 15-ppm precursor ion tolerance and a 0.5 Da fragment ion tolerance, while oxidation of methionine and phosphorylation of serine, threonine and tyrosine were set as variable modifications, whereas cysteine carbamidomethylation was set as fixed modification. A maximum of 5 modifications per peptide was allowed. Trypsin was selected as protease allowing up to two missed cleavages, and peptides were limited to a maximum of 4600 Da with minimum and maximum lengths of 8 and 25 amino acids respectively. PSMs were adjusted to a 1% false discovery rate (FDR), and a minimum of two peptides with at least one unique peptide was required. Match between runs was enabled with a match time window of one minute and an alignment time window of 20 minutes. The type of analysis was set to reporter ion MS2, and reporter ion intensities were adjusted to correct for isotopic impurities of the different TMT 6-plex reagents according to the manufacturer's specifications.

Both neutral loss (NL) MaxQuant phosphopeptides datasets were analyzed separately in R (r-project.org). After removing "Reverse", "Potential Contaminants," and phosphopeptides showing a localization probability of less than 0.75, we applied a modified version of the internal reference scaling (IRS) normalization procedure [7]. First, we applied a median normalization to each TMT 6-plex experiment. We next corrected the unavoidable between-run technical variations caused by the stochastic selection of peptide ions during MS data acquisition. For each phosphopeptide of each of the three TMT experiments, we summed the MS2 reporter ions intensities of the six channels. These reference values were then averaged by geometric mean, allowing us to estimate the scaling factor of each phosphopeptide, and to adjust their reference value to their respective geometric mean. Statistical contrasts were performed with the exactTest function after we estimated the dispersion from the EdgeR package [8].

**Bioinformatics**

We considered as modulated phosphosites those showing a normalized Log2 ($\leq -0.5$ or $\geq 0.5$) between two given conditions and $P < 0.05$. Reactome pathway analyses were performed to functionally annotate proteins using *Gene Set Enrichment Analysis* (*GSEA*) with an FDR threshold of 0.05. Graphical network representations of protein-protein interaction were performed with the STRING app (11.5) in Cytoscape (3.9.1). Volcano plots were generated using GraphPad Prism 10.

**Statistics**

Statistical analyses were performed using GraphPad Prism 10 software. Results are shown as the mean ± SEM. For single measurements, an unpaired, 2-tailed Student's *t* test was used, while 1-way ANOVA followed by Bonferroni's post test was used for comparison of more than 2 groups. For repeated measurements, a repeated-measures 2-way ANOVA followed by Bonferroni's post test was used. In all analysis, $p<0.05$ was considered statistically significant.

**Study Approval**

All animal use complied with the guidelines of the Canadian Committee for Animal Protection and was approved by the IRCM Animal Care Committee.

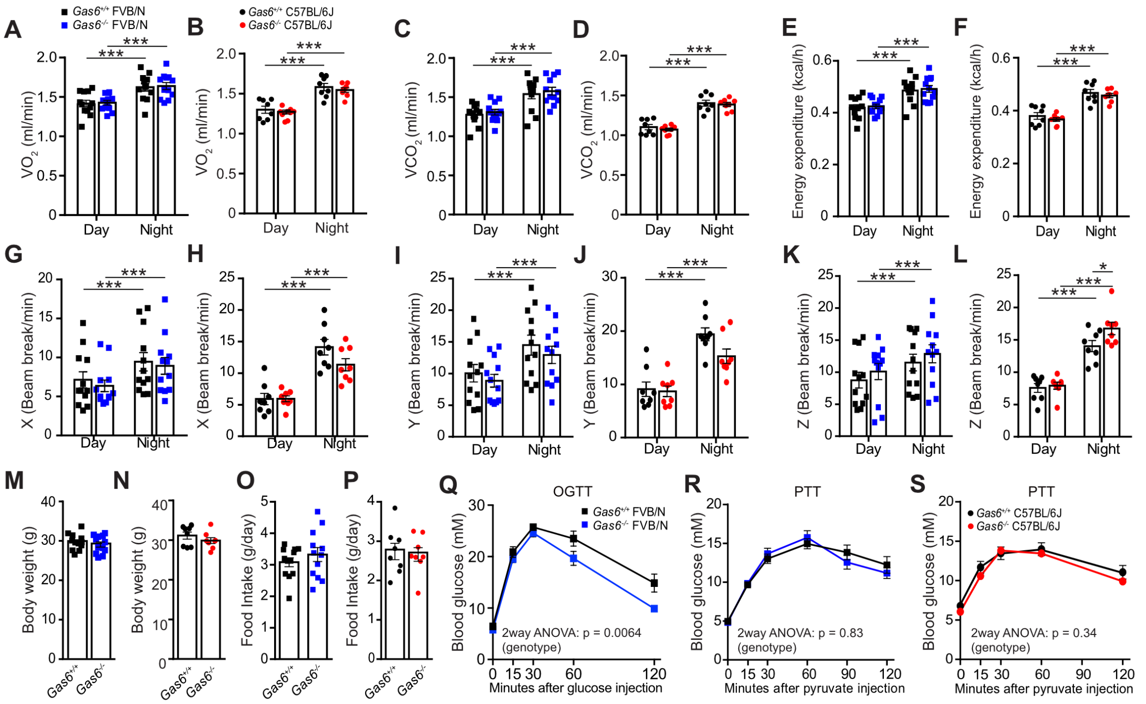


**Figure S1. GAS6 deficiency does not affect energy expenditure, food intake or gluconeogenesis.** Metabolic parameters of 3-months old *Gas6*$^{+/+}$ and *Gas6*$^{-/-}$ FVB/N (blue) and C57BL/6J (red) male mice (n=8-12). **(A-B)** $O_2$ consumption, **(C-D)** $CO_2$ release, **(E-F)** energy expenditure, **(G-H)** ambulatory activity on the X axis, **(I-J)** Y axis and **(K-L)** Z axis. **(M-N)** Body weight. **(O-P)** Food intake. **(Q)** Oral glucose tolerance test (OGTT). Mice were in mited for 16h. Mice received orally a 1.5g/kg bolus of glucose. **(R-S)** Pyruvate tolerance tests (PTT) in mice fasted 16h and i.p. injected with 2g/kg of pyruvate (n=8-17). Results represent the mean ± SEM. **P* < 0.05 and ****P* < 0.001, by two-way ANOVA for repeated measurement followed by Bonferroni's post tests **(A-L and Q-S)** or by unpaired, two-tailed Student's *t* test **(M-P)**.

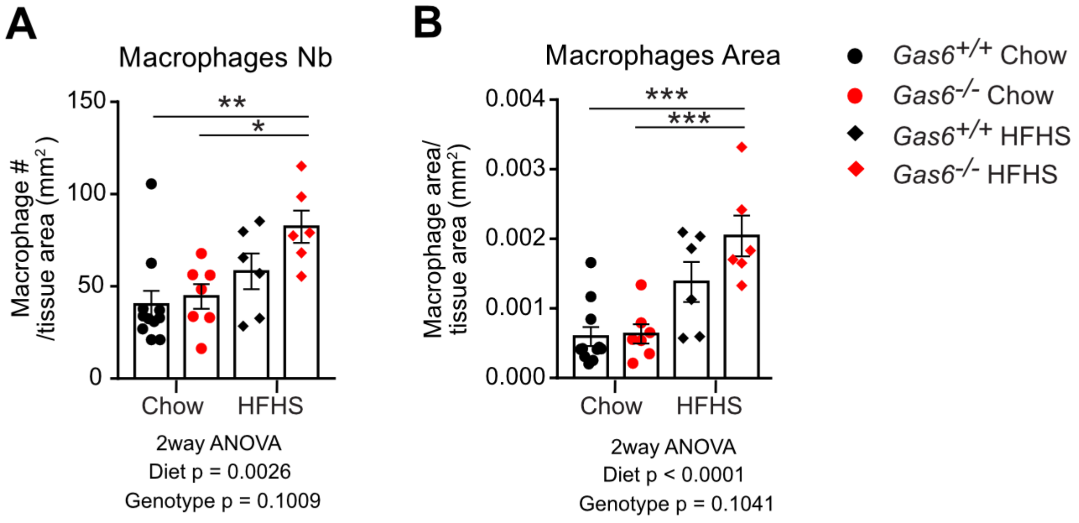


**Figure S2. GAS6 deficiency does not protect against inflammation induced by High-Fat/High-Sucrose diet.** Quantification of macrophages infiltration in epididymal white adipose tissue (WAT) by F4/80 immunohistochemical staining in *Gas6*$^{+/+}$ and *Gas6*$^{-/-}$ mice fed chow diet (Chow) or High-Fat/High-Sucrose diet (HFHS). **(A)** Quantification of F4/80 positive macrophage number in eWAT. **(B)** Quantification of total area of F4/80 positive signal. Results are the average of 5 sections per mouse (n=6-11 mice per genotype and diet) and are normalized over tissue area (mm$^2$). Results represent mean ± SEM, *$P < 0.05$, **$P < 0.01$, ***$P < 0.001$ by two-way ANOVA with Bonferroni's post tests.

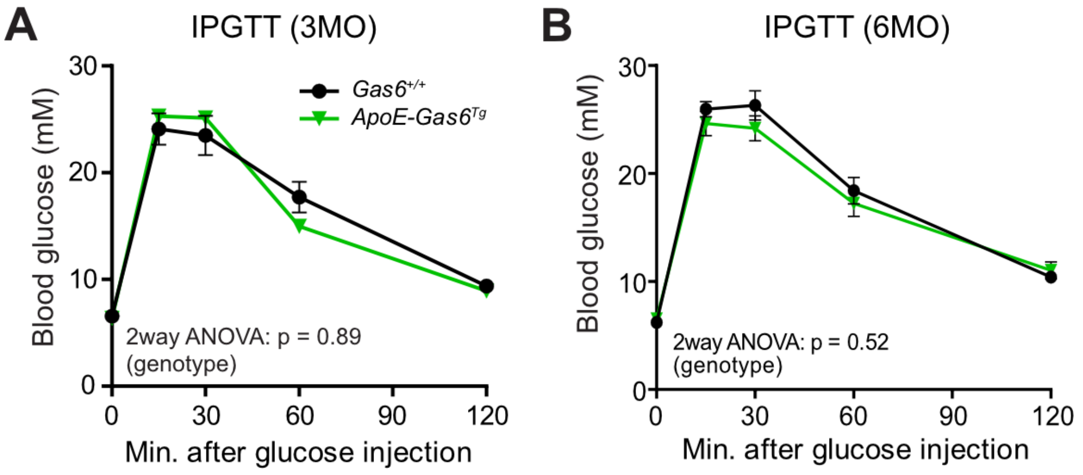


**Figure S3. Glucose tolerance in transgenic *ApoE-Gas6$^{Tg}$* mice. (A-B)** Intra-peritoneal glucose tolerance test (IPGTT) in *Gas6$^{+/+}$* and *ApoE-Gas6$^{Tg}$* mice that were subjected to a 16h-fast and injected i.p. with 2g/kg of glucose at 3 and 6 months of age (n=11-19). Results represent mean ± SEM. Two-way ANOVA for repeated measurements with Bonferroni's post tests revealed no significant difference.

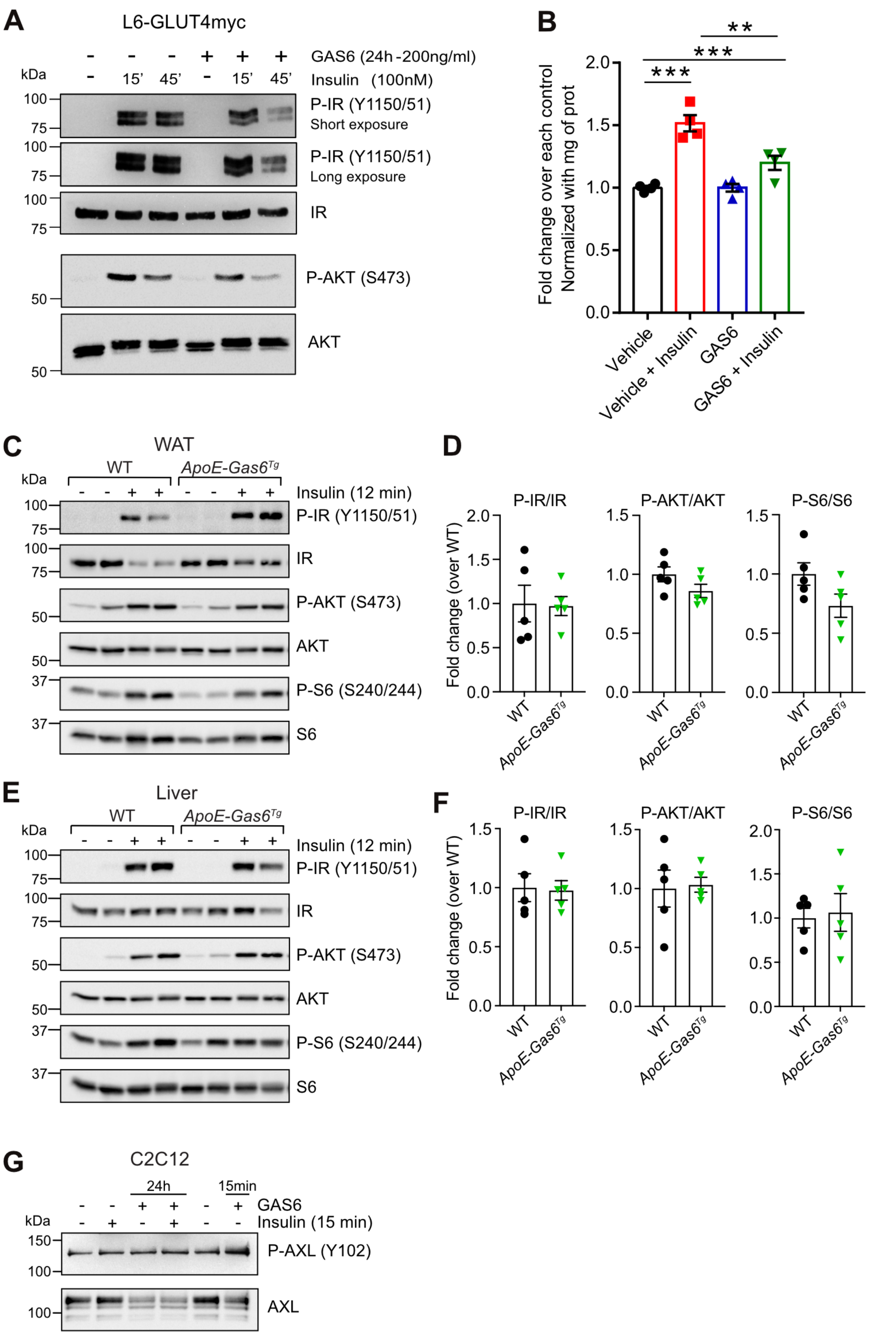
A
L6-GLUT4myc
GAS6 (24h -200ng/ml)
Insulin (100nM)
kDa
15'
45'
100
75
50
P-IR (Y1150/51)
Short exposure
P-IR (Y1150/51)
Long exposure
IR
P-AKT (S473)
AKT
B
Fold change over each control
Normalized with mg of prot
2.0
1.5
1.0
0.5
0.0
***
***
**
Vehicle
Vehicle + Insulin
GAS6
GAS6 + Insulin
C
WAT
WT
ApoE-Gas6^Tg
Insulin (12 min)
kDa
37
P-IR (Y1150/51)
IR
P-AKT (S473)
AKT
P-S6 (S240/244)
S6
D
P-IR/IR
P-AKT/AKT
P-S6/S6
Fold change (over WT)
WT
ApoE-Gas6^Tg
E
Liver
WT
ApoE-Gas6^Tg
Insulin (12 min)
P-IR (Y1150/51)
IR
P-AKT (S473)
AKT
P-S6 (S240/244)
S6
F
P-IR/IR
P-AKT/AKT
P-S6/S6
Fold change (over WT)
WT
ApoE-Gas6^Tg
G
C2C12
24h
15min
GAS6
Insulin (15 min)
kDa
150
100
P-AXL (Y102)
AXL

**Figure S4. GAS6 affects insulin signaling and GLUT4 translocation in muscle cells.** **(A)** Insulin signaling pathway activation in L6-GLUT4myc cells treated with GAS6 (200 ng/ml) for 24h followed by insulin stimulation (100 nM) for 15 or 45 minutes. Phosphorylation of IR (Y1150/1151) and AKT (S473) was assessed by Western blot analysis. Total IR and total AKT were used as loading controls. Representative from 3 independent experiments. **(B)** Surface GLUT4 levels in L6-GLUT4myc myoblasts treated for 24 hours with GAS6 (200ng/ml) and measured in response to stimulatory insulin (100 nM-15min). A colorimetric assay was used to quantify myc-tagged GLUT4 at the cell surface. Results are expressed as fold change of insulin response relative to each corresponding control (Vehicle vs Vehicle + Insulin and GAS6 vs GAS6+ Insulin). **(C)** Phosphorylation of the insulin receptor (Y1150/1151), AKT (S473) and ribosomal protein S6 (S240/244) in white adipose tissue (WAT) of WT and *ApoE-Gas6*$^{Tg}$ mice 12 minutes after an i.v. injection of saline (-) or insulin (0.5U/kg). **(D)** Quantification of the phosphorylation levels normalized over the total amount of each protein (n=5). **(E)** Phosphorylation of the insulin receptor (Y1150/1151), AKT (S473) and ribosomal protein S6 (S240/244) in liver of WT and *ApoE-Gas6*$^{Tg}$ mice 12 minutes after an i.v. injection of saline (-) or insulin (0.5U/kg). **(F)** Quantification of the phosphorylation levels normalized over the total amount of each protein (n=5). **(G)** Phosphorylation of AXL (Y102) was assessed by Western blot in C2C12 cells treated with GAS6 (200 ng/ml) for 24h followed by insulin stimulation (100 nM) for 15 minutes. Total AXL was used as loading control. Results represent mean ± SEM (n=4 per condition). **$P < 0.01$, ***$P < 0.001$, by one-way ANOVA followed by Bonferroni's post tests **(B),** or by unpaired, two-tailed Student's *t* test **(D, F)**.

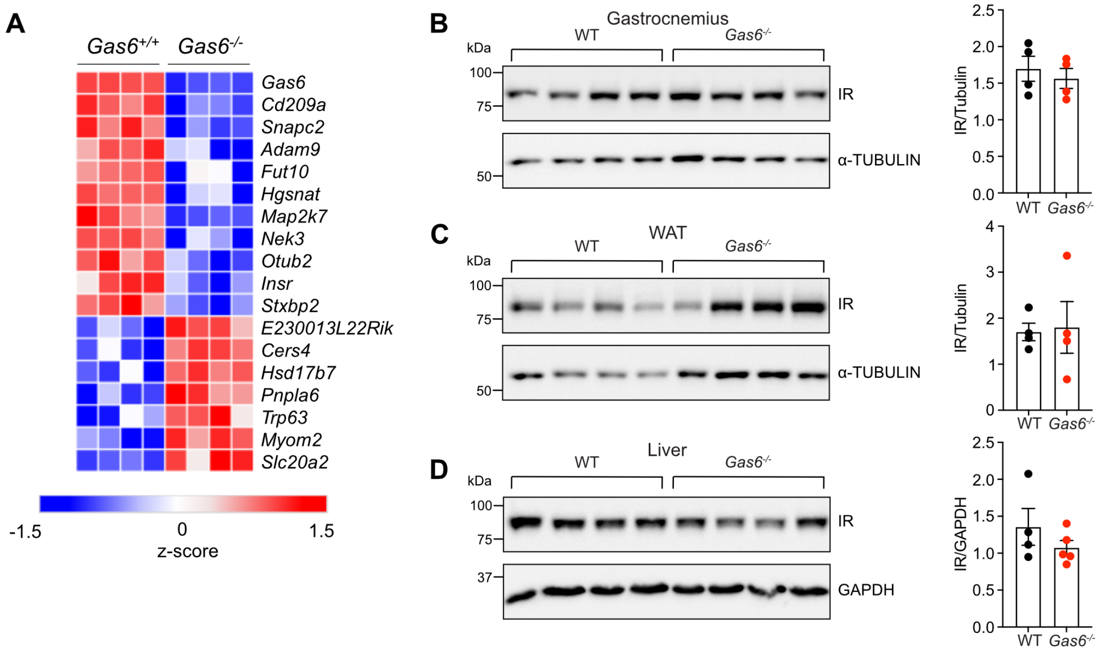


**Figure S5. Transcription profile and Insulin Receptor protein level in skeletal muscle of GAS6-deficient mice.**

**(A)** Heatmap representation of the 18 differentially expressed genes in skeletal muscle (gastrocnemius) of *Gas6*$^{-/-}$ mice (n=4) by RNA-sequencing (p-adj < 0.05). Blue and red represent the significantly downregulated and upregulated genes respectively. **(B-D)** Insulin receptor (IR) protein levels in WT and *Gas6*$^{-/-}$ mice assessed by Western blot in gastrocnemius **(B)**, white adipose tissue (WAT) **(C)** and liver **(D)**. Alpha-tubulin or Gapdh was used as loading control. Results represent mean ± SEM, by unpaired, 2-tailed Student's *t* test **(B-D).**

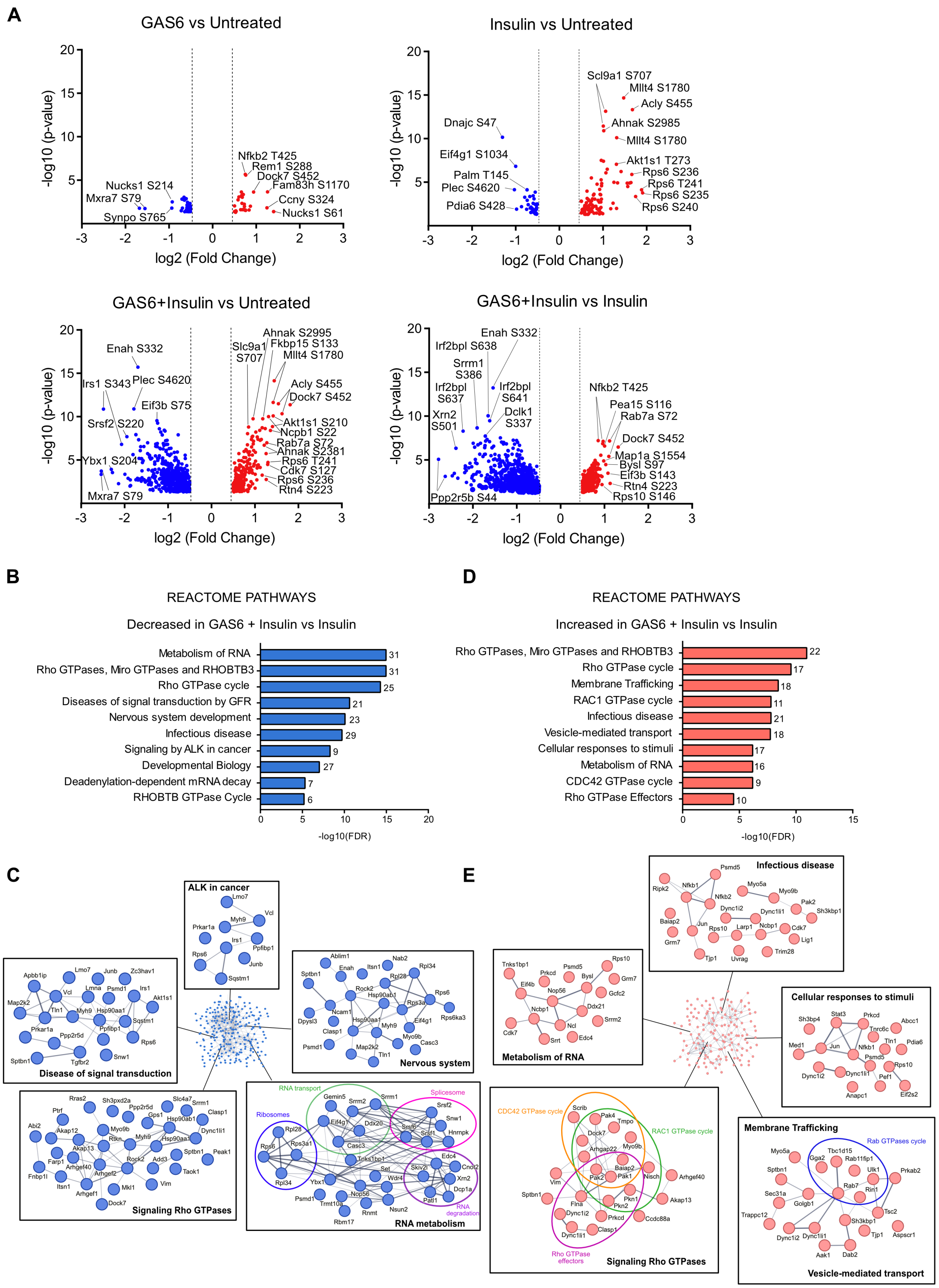
A
GAS6 vs Untreated
Nfkb2 T425
Rem1 S288
Dock7 S452
Fam83h S1170
Ccny S324
Nucks1 S61
Nucks1 S214
Mxra7 S79
Synpo S765
Insulin vs Untreated
Scl9a1 S707
Mllt4 S1780
Acly S455
Dnajc S47
Ahnak S2985
Mllt4 S1780
Eif4g1 S1034
Akt1s1 T273
Rps6 S236
Palm T145
Rps6 T241
Plec S4620
Rps6 S235
Pdia6 S428
Rps6 S240
GAS6+Insulin vs Untreated
Ahnak S2995
Slc9a1 S707
Fkbp15 S133
Mllt4 S1780
Enah S332
Acly S455
Irs1 S343
Plec S4620
Dock7 S452
Eif3b S75
Akt1s1 S210
Srsf2 S220
Ncpb1 S22
Rab7a S72
Ahnak S2381
Ybx1 S204
Rps6 T241
Cdk7 S127
Rps6 S236
Mxra7 S79
Rtn4 S223
GAS6+Insulin vs Insulin
Enah S332
Irf2bpl S638
Srrm1 S386
Irf2bpl S641
Irf2bpl S637
Dclk1 S337
Xrn2 S501
Nfkb2 T425
Pea15 S116
Rab7a S72
Dock7 S452
Map1a S1554
Bysl S97
Eif3b S143
Rtn4 S223
Rps10 S146
Ppp2r5b S44
-log10 (p-value)
log2 (Fold Change)
B
REACTOME PATHWAYS
Decreased in GAS6 + Insulin vs Insulin
Metabolism of RNA 31
Rho GTPases, Miro GTPases and RHOBTB3 31
Rho GTPase cycle 25
Diseases of signal transduction by GFR 21
Nervous system development 23
Infectious disease 29
Signaling by ALK in cancer 9
Developmental Biology 27
Deadenylation-dependent mRNA decay 7
RHOBTB GTPase Cycle 6
-log10(FDR)
D
REACTOME PATHWAYS
Increased in GAS6 + Insulin vs Insulin
Rho GTPases, Miro GTPases and RHOBTB3 22
Rho GTPase cycle 17
Membrane Trafficking 18
RAC1 GTPase cycle 11
Infectious disease 21
Vesicle-mediated transport 18
Cellular responses to stimuli 17
Metabolism of RNA 16
CDC42 GTPase cycle 9
Rho GTPase Effectors 10
-log10(FDR)
C
ALK in cancer
Disease of signal transduction
Nervous system
Signaling Rho GTPases
RNA transport
Spliceosome
Ribosomes
RNA degradation
RNA metabolism
E
Infectious disease
Metabolism of RNA
Cellular responses to stimuli
CDC42 GTPase cycle
RAC1 GTPase cycle
Rho GTPase effectors
Signaling Rho GTPases
Membrane Trafficking
Rab GTPases cycle
Vesicle-mediated transport

**Figure S6. GAS6 rewires insulin signaling pathways in muscle cells. (A)** Volcanoplots of phosphosites significantly changed ($P < 0.05$) between the different treatments. Fold change is represented as and log2 Fold change $\geq 0.5$ (red plots) or $\leq -0.5$ (blue plots). **(B)** Reactome pathway analysis of proteins with decreased phosphorylation when C2C12 cells were treated with GAS6 in combination with insulin compared to insulin alone, generated by Gene Set Enrichment Analysis (GSEA) (FDR q-value < 0.05). The number of proteins associated to each pathway is indicated next to each bar graphs. **(C)** Protein-protein interaction network analysis of proteins with decreased phosphorylation when treated with GAS6 and insulin compared to insulin alone, generated using STRING and Cytoscape. **(D)** Results of GSEA Reactome pathway analysis of proteins with increased phosphorylation when treated with GAS6 in combination with insulin compared to insulin alone (FDR q-value < 0.05). The number of proteins associated to each pathway is indicated next to each bar graphs. **(E)** Network layout of protein-protein interaction of proteins with increased phosphorylation when C2C12 cells were treated with GAS6 in combination with insulin compared to insulin alone, generated using STRING and Cytoscape.

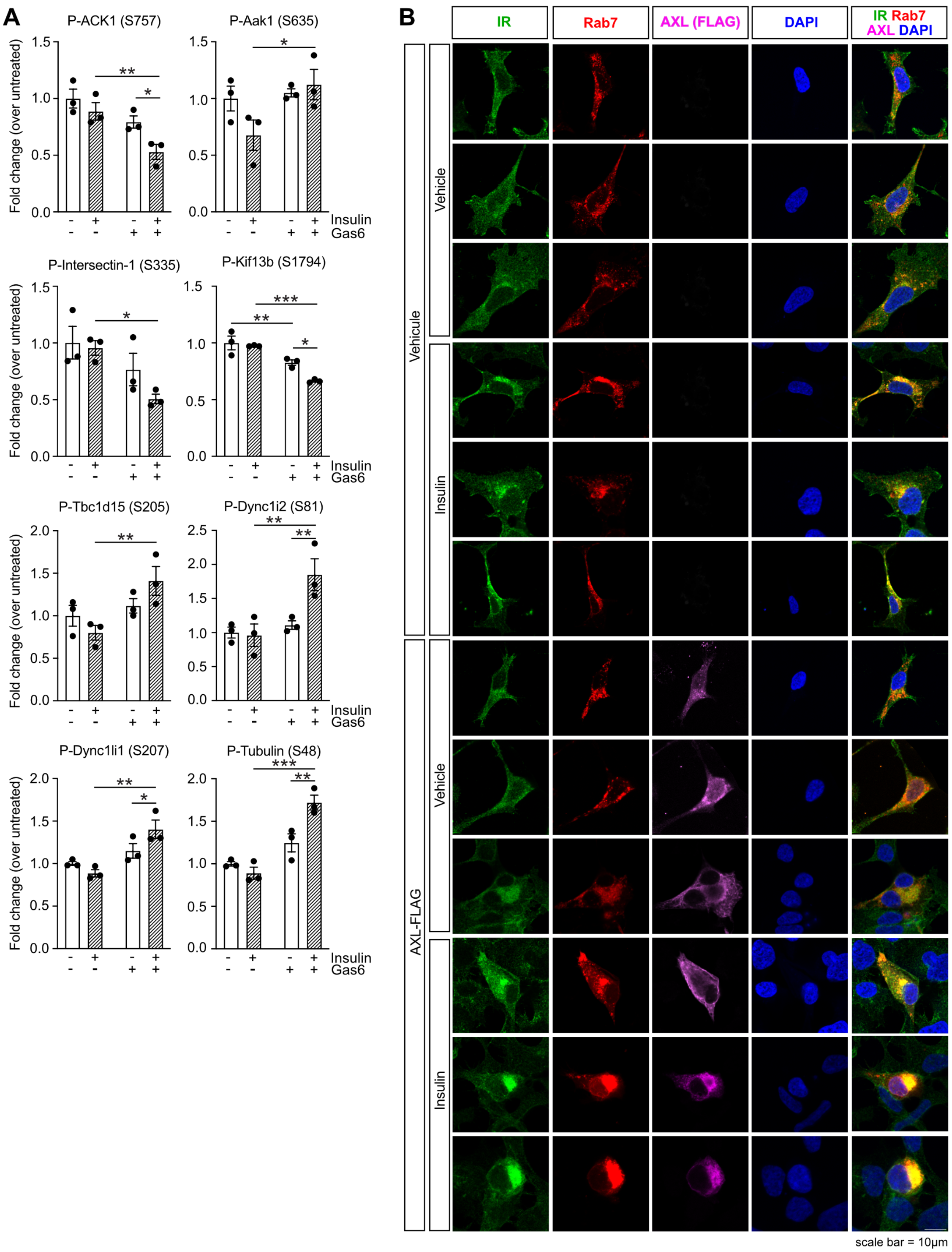
A
P-ACK1 (S757)
P-Aak1 (S635)
P-Intersectin-1 (S335)
P-Kif13b (S1794)
P-Tbc1d15 (S205)
P-Dync1i2 (S81)
P-Dync1li1 (S207)
P-Tubulin (S48)
Fold change (over untreated)
Insulin
Gas6
B
IR
Rab7
AXL (FLAG)
DAPI
IR Rab7 AXL DAPI
Vehicule
Vehicle
Insulin
AXL-FLAG
scale bar = 10µm

**Figure S7. (A)** Phosphorylation levels of proteins measured in the quantitative phosphoproteomics analysis. **(B)** Additional representative confocal microscopy images of HEK293 cells transfected with Rab7-mScarlet and AXL-Flag or empty vector, and treated or not with 100 nM insulin for 15 minutes and quantified in Figure 7C-E. Cells were stained for AXL-Flag (Alexa Fluor 633-magenta) and endogenous IR (Alexa-Fluor 488-green). DAPI was used to stain nuclei. Scale bar, 10 µm. Related to Figure 7C. Results represent mean ± SEM (n=3). *$P < 0.05$, **$P < 0.01$, ***$P < 0.001$ by 2-way ANOVA followed by Fisher's LSD post test.

### Supplemental Table 1

| Gene | Sequence 5'-3' | Purpose |
|---|---|---|
| *Axl-Fw* | ATAGGGCTAACTCGAGAGG | qPCR |
| *Axl-Rv* | GGCTCTAGGGGCACAGGAAG | qPCR |
| *Gas6-Fw* | GATACGGCCTCGTACAAGCA | qPCR |
| *Gas6-Rv* | CTCCAGTGTTGCAGGACCAT | qPCR |
| *Mertk-Fw* | CAGTTGCTAGAGAGCTGCGA | qPCR |
| *Mertk-Rv* | GGTGTGACTGCAGCAAAAGG | qPCR |
| *Tyro3-Fw* | TGTCTGCGAATGGAACTGGA | qPCR |
| *Tyro3-Rv* | TGCCCTGGTGACTCGGATAG | qPCR |
| *S16-Fw* | AGGAGCGATTTGCTGGTGTGG | qPCR |
| *S16-Rv* | GCTACCAGGGCCTTTGAGA | qPCR |
| *Gas6-Fw* | GAGTGCCGTGATTCTGGTC | Genotyping-*Gas6* |
| *Gas6-Fw* | ATCTCTCGTGGGATCATT | Genotyping-Gas6 |
| *Gas6-Rv* | CCACTAAGGAAACAATAACTG | Genotyping-*Gas6* |
| *ApoE prom-Fw* | AAGGCTAACCTGGGGTGAGG | Genotyping-*ApoE-Gas6Tg* |
| *Gas6-Rv* | AAGTTCTGAACACATTTGGCGA | Genotyping-*ApoE-Gas6Tg* |
| *IL-2-Fw* | CTAGGCCACAGAATTGAAAGATCT | Genotyping |
| *IL-2-Rv* | GTAGGTGGAAATTCTAGCATCATCC | Genotyping |
| *IR-Fw* | TTAAAAGCTTGCTCTGATCCGAGGAGACCC | Cloning |
| *IR-Rv* | ATTGTCGACGGAAGGATTGGACCGAGG | Cloning |
| HA-tag-Fw | P- TCGACTACCCATACGATGTTCCAGATTACGCTTAAG | Cloning |
| HA-tag-Rv | P- AATTCTTAAGCGTAATCTGGAACATCGTATGGGTAG | Cloning |
| *Gas6-Fw* | ATTACAAGCTTGCCACCATGCCGCCACCG | Cloning |
| *Gas6-Rv* | TTAATCTAGAGGGGTGGCATGCTCCACAGG | Cloning |
| *Rab7a-Fw* | CATCATAAGCTTCGATGACCTCTAGGAAGAAAGTG | Cloning |
| *Rab7a-Rv* | CATCATGGGCCCTTAACAACTGCAGCTTTCTG | Cloning |

### Supplemental Table 2

| Antibody | Company | Catalog # | Application | Dilution |
|---|---|---|---|---|
| Gamma-carboxyglutamyl (Gla) residues | Sekisui Diagnostics | 3570 | WB | 1/500 |
| Myc-Tag (9B11) | Cell Signaling | 2276 | WB | 1/1000 |
| phospho-AXL (Y702) | Gene Script (custom made) | N/A | WB | 1/1000 |
| Axl (C-20) | SantaCruz | sc-1096 | WB | 1/500 |
| phospho-AKT (Ser473) | Cell Signaling | 9271 | WB | 1/1000 |
| AKT (pan) (C67E7) | Cell Signaling | 4691 | WB | 1/1000 |
| phospho-IGF-IR β (Tyr1135/1136)/IR β (Tyr1150/1151) | Cell Signaling | 3024 | WB | 1/1000 |
| Insulin Receptor β (L55B10) | Cell Signaling | 3020 | WB | 1/1000 |
| S6 Ribosomal Protein (5G10) | Cell Signaling | 2217 | WB | 1/1000 |
| phospho-S6 Ribosomal Protein (Ser240/244) | Cell Signaling | 2215 | WB | 1/1000 |
| HA-Tag (C29F4) | Cell Signaling | 3724 | WB | 1/1000 |
| DYKDDDDK Tag (D6W5B) / Flag | Cell Signaling | 14793 | WB | 1/1000 |
| Anti-Mouse HRP | Jackson ImmunoResearch | 115-035-174 | WB | 1/5000 |
| Anti-Rabbit HRP | Jackson ImmunoResearch | 211-032-171 | WB | 1/5000 |
| Monoclonal Anti-Flag ® M2 | Sigma | F1804 | IF | 1/2000 |
| Insulin Receptor β (E9L5V) | Cell Signaling | 23413 | IF | 1/100-1/200 |
| Alexa Fluor 488-Donkey Anti-rabbit IgG | Jackson ImmunoResearch | 711-545-152 | IF | 1/1000 |
| Cy3-Donkey Anti-Mouse IgG | Jackson ImmunoResearch | 715-165-150 | IF | 1/500 |
| Alexa Fluor 633 Goat Anti-mouse IgG | Invitrogen | A21050 | IF | 1/500 |
| Laminin DyLight 650 | Novus Biochemicals | NB300-144C | IF | 1/200 |
| F4/80 | Cell Signaling | 70076 | IHC | 1/500 |

WB: western blot; IF: Immunofluorescence; IHC: Immunohistochemistry
